# Accurate Identification and Mechanistic Evaluation of Pathogenic Missense Variants with *Rhapsody-2*

**DOI:** 10.1101/2025.02.17.638727

**Authors:** Anupam Banerjee, Anthony Bogetti, Ivet Bahar

**Affiliations:** Laufer Center for Physical and Quantitative Biology, Stony Brook University, New York 11794, USA; Department of Biochemistry and Cell Biology, Renaissance School of Medicine, Stony Brook University, New York 11794, USA

**Author notes:** Ivet Bahar^1,2^ and Anupam Banerjee^1^, **Email:**.

**Keywords:** Pathogenicity Prediction, Structural Dynamics, Missense Variants, Machine Learning

## Abstract

Understanding the effects of missense mutations or single amino acid variants (SAVs) on protein function is crucial for elucidating the molecular basis of diseases/disorders and designing rational therapies. We introduce here Rhapsody-2, a machine learning tool for discriminating pathogenic and neutral SAVs, significantly expanding on a precursor limited by the availability of structural data. With the advent of AlphaFold2 as a powerful tool for structure prediction, Rhapsody-2 is trained on a significantly expanded dataset of 117,525 SAVs corresponding to 12,094 human proteins reported in the ClinVar database. Adopting a broad set of descriptors composed of sequence evolutionary, structural, dynamic, and energetics features in the training algorithm, Rhapsody-2 achieved an AUROC of 0.94 in 10-fold cross-validation when all SAVs of a particular test protein (mutant) were excluded from the training set. Benchmarking against a variety of testing datasets demonstrated the high performance of Rhapsody-2. While sequence evolutionary descriptors play a dominant role in pathogenicity prediction, those based on structural dynamics provide a mechanistic interpretation. Notably, residues involved in allosteric communication, and those distinguished by pronounced fluctuations in the high frequency modes of motion or subject to spatial constraints in soft modes usually give rise to pathogenicity when mutated. Overall, Rhapsody-2 provides an efficient and transparent tool for accurately predicting the pathogenicity of SAVs and unraveling the mechanistic basis of the observed behavior, thus advancing our understanding of genotype-to-phenotype relations.

**Significance Statement:** Understanding the impact of single amino acid variants (SAVs) on protein function is pivotal for detecting and treating genetic disorders. We present *Rhapsody-2*, a machine learning tool that leverages AlphaFold2-predicted protein structures to accurately predict the pathogenicity of SAVs. By integrating sequence, structure, and dynamics properties, *Rhapsody-2* achieves a high performance in distinguishing pathogenic variants across multiple benchmark datasets. Our tool not only provides a robust framework for predicting the impact of SAVs but also allows for mechanistic interpretability of pathogenicity, offering a valuable resource for understanding genotype-to-phenotype relationships toward assisting in precision medicine.

## Introduction

Missense mutations are genetic variations in which one or more nucleotide changes in the DNA lead to amino acid substitutions in the encoded protein. Unlike silent mutations, missense mutations can have a range of effects on the structure, function, and interactions of the encoded single amino acid variant (SAV). Pathogenic SAVs may exhibit severe disruptions in folding, enzymatic and other activities, and interactions with other proteins. These disruptions manifest as genetic diseases, cancer, and neurological disorders (1-4).

The human exome harbors a large number (76 million) of potential missense variants (5). Accurate prediction of the effects, pathogenic or benign, of these variants is a primary goal across clinical practice and public health. Such predictions may guide appropriate diagnosis and treatment strategies and facilitate disease prognosis and risk assessments, provided that they are made in the context of sequence evolution, structure, dynamics, and interactions of the mutants in the cell.

In recent years, many computational approaches powered by advances in machine or deep learning (ML or DL) and artificial intelligence (AI) techniques, as well as those employing robust traditional methods, have been developed for SAV pathogenicity predictions. These approaches include Missense3D (6), a homology modeling-aided structure-based predictor, SIFT (7), which uses sequence homology to compute the likelihood that an amino acid substitution will have an adverse effect on protein function; REVEL (8), an ensemble method that combines the results of multiple methods; EVE (9), a deep, generative model of evolutionary data; LYRUS (10), an ML predictor based on sequence, structure, and dynamics features; SPRI (11), a structure-based pathogenicity relationship identifier; PolyPhen-2 (12), an ML classifier enabled by high-quality multiple sequence alignments; and EVMutation (13), which leverages coevolution information to predict the fitness of mutants; WS-SNPs&GO (14), a server that uses sequence, structure, and GO annotations to predict diseases associated with SNPs; and MutPred (15), an early predictor of pathogenicity, incorporated both functional and structural properties, such as catalytic activity and post-translational modifications, to estimate disease mechanisms. The updated MutPred2 (16) includes a wide range of properties, including secondary structure, signal peptides, and transmembrane topology. MVP and ESM1b are state-of-the-art tools for predicting missense variant pathogenicity. MVP uses a supervised deep residual network trained on labeled pathogenic and benign variants, integrating evolutionary, structural, and gene-specific features (17). In contrast, ESM1b, an unsupervised protein language model, predicts effects across ∼450 million missense variants using sequence-based representations, excelling in isoform-specific and complex variant analyses like in-frame indels (18).

A recent addition to this arsenal is AlphaMissense (19), an *ab initio* deep-learning method, relying on evolution information, which fine-tunes AlphaFold (20) predictions of human and primate variant population frequency databases. Of more than 71 million missense variants in the human proteome, 32% are classified by AlphaMissense as pathogenic, 57% as benign, and 11% as variants of uncertain significance (19). AlphaMissense integrates sequence and structural context by combining co-evolutionary insights from multiple sequence alignments with AlphaFold-derived structural embeddings. It models sequence context using unsupervised protein language modeling and masked residue prediction to capture amino acid distributions and evolutionary constraints. Structural context is encoded through AlphaFold’s pair representations and positional embeddings, processed via Evoformer (20) layers to align sequence and structure features. Fine-tuning on population frequency data enables robust pathogenicity predictions across the proteome, including rare or novel variants, while maintaining high precision in both structured and disordered regions.

These tools have driven remarkable advances in the field. Yet, accurate prediction of pathogenicity by itself is not sufficient for understanding the mechanistic basis of observed behavior, nor does it provide insights into the design of rational intervention strategies. As recently pointed out (21), the merger of ML/AI methods with those based on physical sciences holds great promise for gaining a deeper understanding of target proteins’ structural dynamics at multiple scales; and structural dynamics, in turn, defines the mechanisms of motions accessible to achieve biological function and interactions, whose disruption (due to point mutations at critical sites) may cause dysfunction. The need for considering dynamic effects to understand pathogenicity may extend to tissue-specific non-Mendelian mutations, as suggested by the examination of EGFR ectodomain mutations on kinase domain activation (22). A combined approach that mutually benefits from advances in ML/AI methodology and fundamental theory and concepts of structural biophysics can help provide accurate estimates of pathogenicity *and* shed new insights into the molecular basis of ML/AI predictions, thus potentially accelerating the discovery of intervention methods. The current study aims at achieving those two goals.

The integrated approach between the two disciplines (ML/AI and biophysics) needs a biophysical model and method that lends itself to high throughput generation of data (at the proteome scale), which may then be used for training ML models. Elastic network models (ENMs) meet this requirement. The simplicity of ENMs allows generating mathematically exact analytical solutions, uniquely defined by the protein architecture, for the collective modes of motion intrinsically accessible to the structure, which often underlie biological function.

In 2018, we developed a dynamics-dependent ML-based predictor of SAV pathogenicity (23) and implemented it in the interface *Rhapsody* (24). *Rhapsody* used as features not only sequence and structure data, but also ENM-based descriptors of protein dynamics. Strikingly, despite the simplicity of the approach (a random forest algorithm with only ten features, including three sequence-dependent, one structure-dependent and four dynamics-dependent), the method, benchmarked against a dataset of about 20,000 annotated variants, outperformed the (then) state-of-the-art methods for classifying SAVs as deleterious/pathogenic or neutral/benign. Given that the major difference with respect to existing tools was the incorporation of protein dynamics, we emphasized that the latter was an important determinant of the impact that missense mutations have on protein function. Since then, *Rhapsody* has been widely used as a sequence-, structure- and dynamics-dependent predictor of pathogenicity, with sequence-specific properties being mainly provided by PolyPhen-2 (12).

Yet, the application of *Rhapsody* depends on the availability of a known structure for the protein of interest. This requirement limited the size of the training set as well as the application to structurally unresolved proteins. Precisely, despite identifying an integrated dataset of 87,726 SAVs by combining five publicly available datasets (25-29), only 27,655 SAVs could be matched with PDB structures (23), and *Rhapsody* was trained on an even smaller dataset of 20,361 SAVs corresponding to 2,423 unique human proteins (24) after selecting the SAVs with sufficiently high-confidence labels in ClinVar (2).

With the availability of structural data for most human proteins made possible by AlphaFold2 (20), we are in a position to address the above limitation. By leveraging the available wealth of structural data accessible in the AlphaFold database, as well as recent advances in ENM methodology and ML algorithms, we present here *Rhapsody-2*, a predictor with significantly higher accuracy and coverage than its predecessor. Importantly, *Rhapsody-2* also provides insights into the mechanistic basis of the predictions. The new training database (*Rhapsody-2* DB) comprises 117,525 SAVs spanning 12,094 human proteins; and the new algorithm considers 100 descriptors, composed of 22 evolutionary, 17 structural, 21 dynamics-based, 33 energetics-based, and 6 physicochemical (residue-specific) features, and one on intermolecular interactions (Supplemental Information (*SI*) **Table S1**).

Accurate evaluation of pathogenicity prediction tools requires addressing biases that occur when variants from the same protein are included in both training and testing datasets, leading to somewhat higher performance (30). To avoid this, *Rhapsody-2* was evaluated using protein-stratified cross-validations or tests. Protein-stratification refers to the inclusion of the SAVs of a particular protein in either the training or the testing dataset, thus preventing data leakage that may originate from a particular protein spanning training and testing datasets. Under these conditions, *Rhapsody-2* achieved an AUROC of 0.94 in 10-fold cross-validations on the *Rhapsody-2* DB, and performances comparable to state-of-the-art tools when tested on independent benchmark datasets. These results address our first of two main objectives of delivering a reliable pathogenicity predictor. Our second objective—understanding the biophysical basis of pathogenicity—is met by identifying the features that differentiate between deleterious and neutral mutations. Overall, *Rhapsody-2* provides both accurate predictions of single amino acid variant (SAV) pathogenicity and insights into their molecular mechanisms, advancing our understanding of genotype-phenotype relationships.

## Results

### *Rhapsody-2* database, features, and methodology

Figure 1. provides a schematic description of the methodology. Among the five categories of features used in *Rhapsody-2*, structural features measure how the mutation is accommodated in the folded structure, including the overall packing and interactions with spatial neighbors. Physicochemical features refer to the specific properties of amino acids in the wild-type (WT) protein and its SAV. Dynamics features refer to the spectrum of equilibrium (collective) motions and allosteric communication properties shortly called intrinsic dynamics, evaluated using two elastic network models (ENMs) implemented in the *ProDy* interface (31, 32): the Gaussian network model (GNM) (33) and the anisotropic network model (ANM) (34, 35). Evolutionary features include sequence conservation and the potential tolerance to single or compensating amino acid variations across sequential homologs using the DIAMOND package (36). Additionally, we assessed the energetic properties of the folded state for both the WT and mutant protein. Detailed descriptions of these attributes are provided in the *SI* Methods.

**Figure 1.**
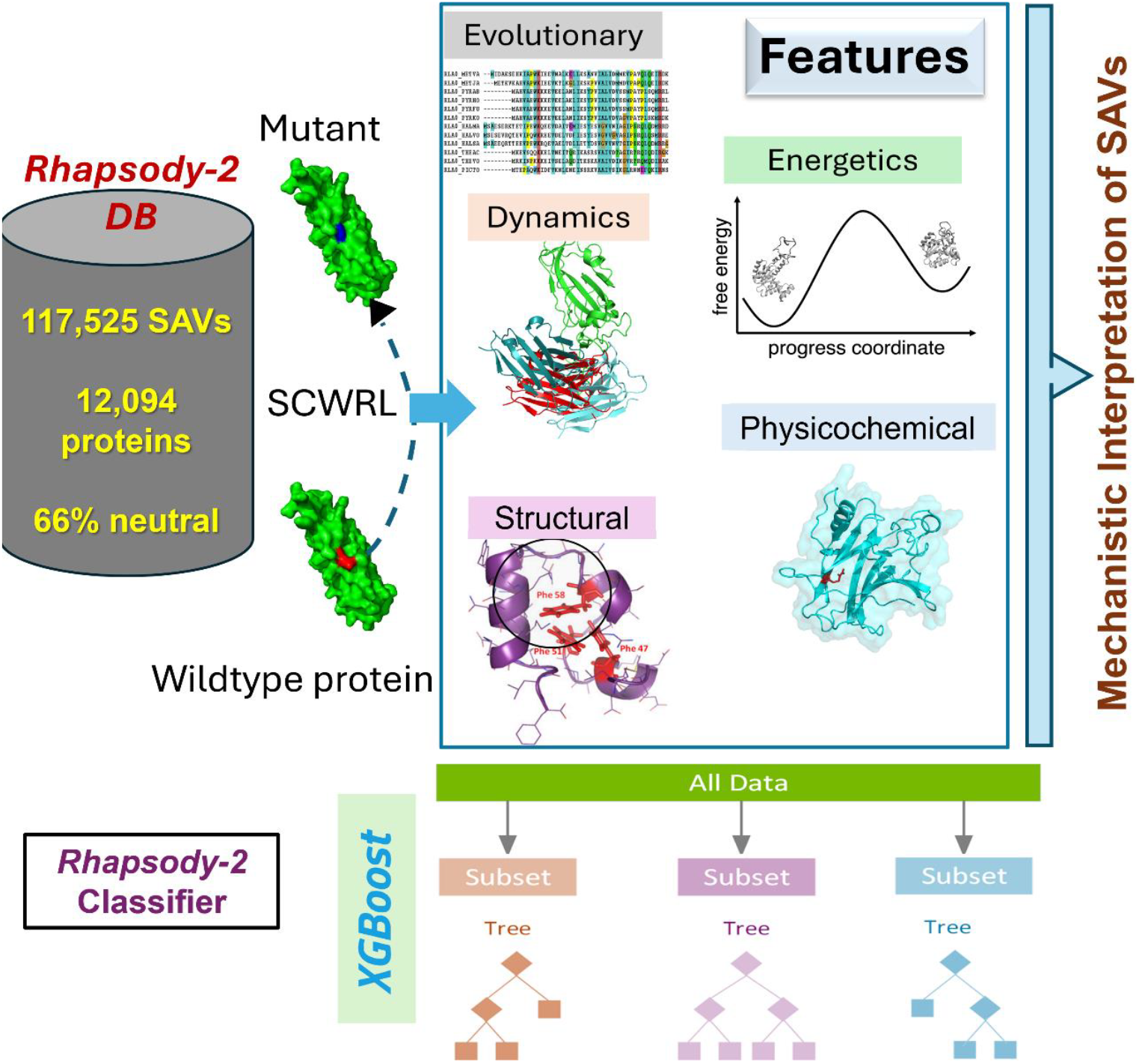
Schematic description of *Rhapsody-2* inputs, method, and outputs. For each SAV in *Rhapsody-2* DB, 100 descriptors composed of 6 residue physicochemical properties, 17 accounting for structural properties, 22 based on sequence evolution, 33 based on energetics, 21 based on intrinsic dynamics, and 1 based on intermolecular interactions were computed. Complete or partial sets of descriptors were used in training *Rhapsody-2* classifier or its variants, respectively, with XGBoost algorithm. Each point mutation was modeled using SCWRL. The distributions of descriptor values in pathogenic and neutral SAVs were used to make inferences on the mechanistic basis of predictions.

Gradient boosting is a machine learning technique that builds models iteratively by adding decision trees, with each tree correcting the errors of the previous ones. XGBoost (eXtreme Gradient Boosting) enhances this method with second-order gradient optimization for precise loss minimization, L1/L2 regularization to prevent overfitting, sparsity-aware algorithms for missing data, and efficient tree pruning for better generalization (37). Its scalability and parallel execution make it highly effective for high-dimensional, structured datasets. We used the XGBoost framework (37) to construct different variants of the *Rhapsody-2* classifier trained on all or a subset of features, including one model exclusively based on intrinsic dynamics.

### Evaluation of the performance of *Rhapsody-2*

Recognizing the potential bias toward accurate prediction when different variants of the same protein appear in both training and testing datasets, as highlighted by Grimm et al. (30), we evaluated *Rhapsody-2* performance averaged over 10-fold protein-stratified cross-validations on the Rhapsody-2 DB. We evaluated the performance based on several metrics: accuracy, precision, recall, F1-score, AUROC and AUPRC (see definitions in *SI* Methods). All metrics, except AUROC and AUPRC, are based on a classification threshold of 0.5 for pathogenicity probability. As shown in **Figure 2A**, *Rhapsody-2* achieved an AUROC of 0.94 and an AUPRC of 0.89 using 10-fold protein-stratified cross-validation. We note that in the absence of protein stratification (**Figure S1**), higher AUROC and AUPRC values of 0.97 and 0.94 are obtained, respectively, showing how the simultaneous inclusion of the same protein’s SAVs in both training and testing datasets overestimates the performance levels.

**Figure 2.**
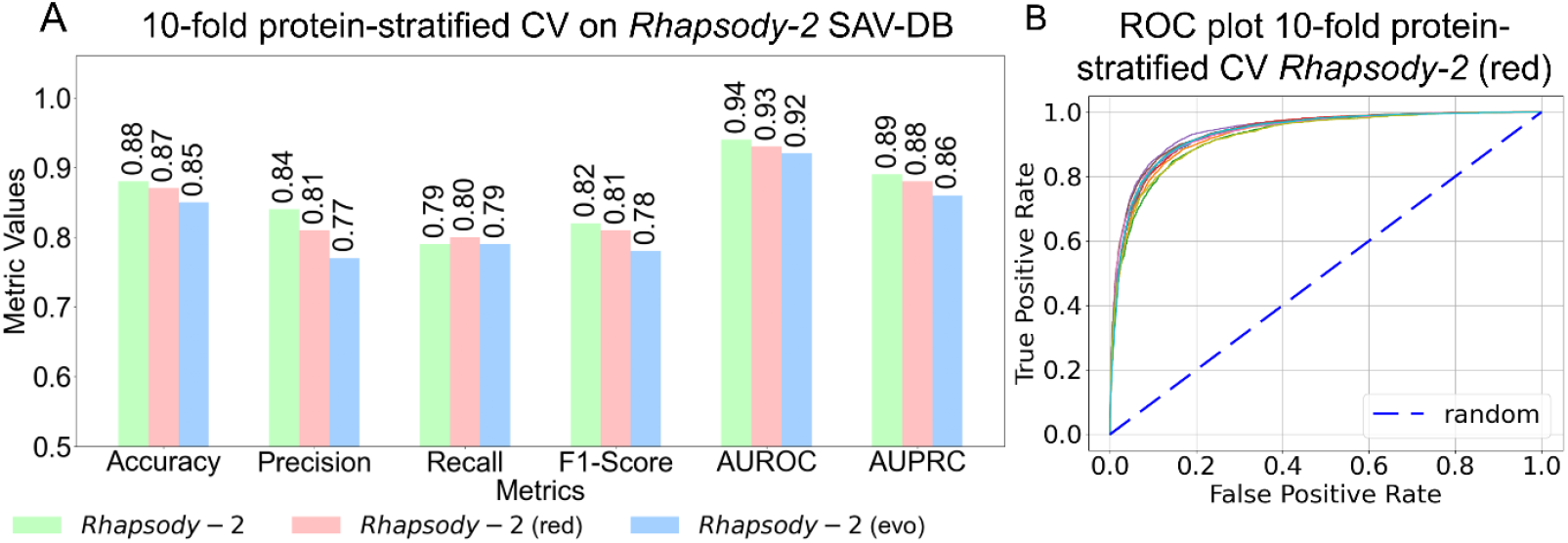
Performance of *Rhapsody-2* and its variants under protein-stratified 10-fold cross-validation on *Rhapsody-2* DB. **(A)** Results based on different metrics (abscissa) for full model (*Rhapsody-2*) and its variants, *Rhapsody-2(red)* and *Rhapsody-2(evo)*, are presented by color-coded bars (see color code at the bottom). **(B)** ROC curves for the reduced model *Rhapsody-2(red)* evaluated under 10-fold protein-stratified cross-validation. The *dashed line* indicates the random behavior.

### Performance of *Rhapsody-2* variants trained on subsets of descriptors

We further examined the performance of *Rhapsody-2* variants, trained on selected subsets of descriptors to assess the contribution of different categories of descriptors to pathogenicity prediction. The *Rhapsody-2(struct)* variant trained exclusively on structural descriptors (listed in rows 77-93 of *SI* **Table S1**) exhibited a mean AUROC of 0.87 under protein-stratified 10-fold cross-validation. *Rhapsody-2*(*dyn*), a model trained solely on dynamics descriptors (rows 23–43 of **Table S1**), achieved an AUROC of 0.79. Note that the dynamics descriptors are based on ENM analyses which exclusively refer to the structure-encoded intrinsic dynamics of the WT protein. These descriptors are agnostic to amino acid identity and are fully defined by the WT inter-residue contact topology. A model trained using sequence evolutionary descriptors only (rows 1-22 in **Table S1**), *Rhapsody-2(evo)*, achieved an AUROC of 0.92 and AUPRC of 0.86 under 10-fold protein stratified CV, confirming that sequence evolutionary information is highly predictive, as also shown by other methods such as *EVE*.

These experiments show that alternative models with comparable performances can be built with different categories of descriptors (each containing approximately 20 features). They further indicate that the different categories of descriptors carry interdependent or partially redundant information. This is consistent with the correlation between sequence evolution and structural dynamics pointed out earlier (38). However, the performance of these variants remains below that of *Rhapsody-2*. Next we explored whether a *reduced model* could be constructed with a limited set of descriptors without significantly compromising on the classification ability of the model.

### Construction of *Rhapsody-2(red)*, a reduced model trained on a small set of descriptors

Our *in silico* experiments in search of a reduced model led to *Rhapsody-2(red)*, trained on two categories of descriptors, evolutionary and dynamics, plus one structural feature, the relative solvent accessibility (RSA). Despite the use of a reduced set of 44 descriptors, this model exhibited a performance almost as strong as the full *Rhapsody-2* (see the *green* and *pink bars* in **Figure 2A**), and even surpassed *Rhapsody-2* in terms of its recall obtained by protein-stratified CV. **Figure 2B** present the ROC plots for *Rhapsody-2(red)* under 10-fold protein stratified CV. *Rhapsody-2(red)* is therefore proposed as a simple and computationally efficient tool: with precomputed data on ENM-based dynamics and evolutionary descriptors, and RSA available for almost all human proteins, it generates *in silico* saturation mutagenesis maps within seconds.

### Benchmarking of *Rhapsody-2* against state-of-the-art datasets

First, we considered the datasets UnifyPDBFull and UnifyPDBAcceptable, introduced alongside the SPRI method (11). We trained *Rhapsody-2* and its variants on these two datasets (using the XGBoost algorithm) and performed 5-fold protein-stratified cross-validations. Panels **A** and **B** in **Figure 3** display the results against the respective UnifyPDBFull and the UnifyPDBAcceptable datasets. *Rhapsody-2* yielded an average AUROC of 0.93 and MCC of 0.70 in panel **A**, closely followed by *Rhapsody-2*(*red*) with an AUROC of 0.92 and an MCC of 0.69. Similarly, on the UnifyPDBAcceptable dataset (panel **B**), *Rhapsody-2* achieved an AUROC of 0.92 and an MCC of 0.67, closely followed by *Rhapsody-2(red)*.

**Figure 3.**
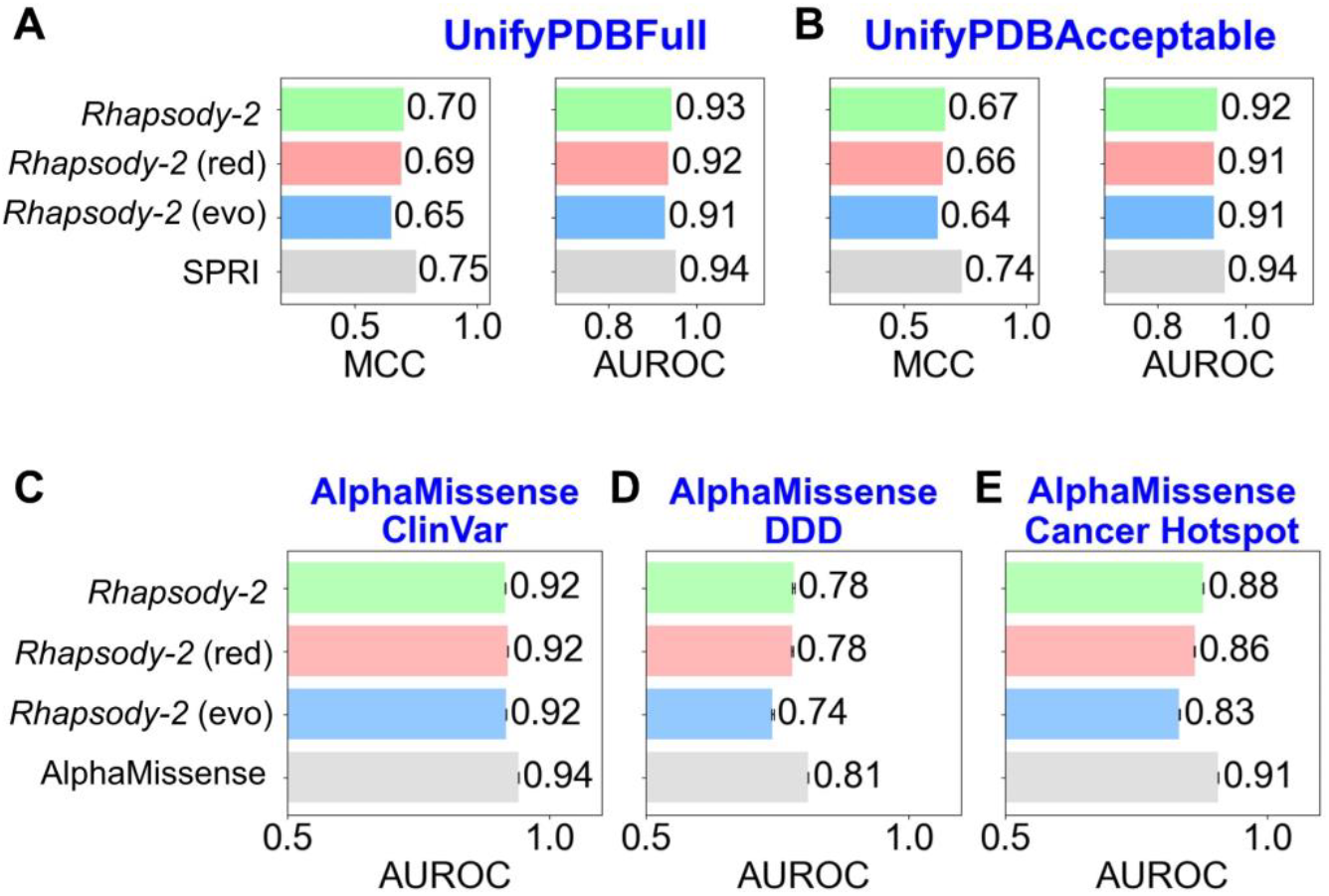
Performance of *Rhapsody-2* and its variants benchmarked against five datasets using protein-stratified tests. **(A-B)** Results from protein-stratified 5-fold cross-validations on UnifyPDBFull (**A**) and UnifyPDBAcceptable (**B**) datasets. **(C-E)** Results from benchmarking against ClinVar, DDD, and Cancer Hotspot databases, respectively. The results from the best-performing methods on these datasets, SPRI in **A-B** and AlphaMissense in **C-E**, are also included for comparison. Note that SPRI and AlphaMissense results were obtained without protein-stratification.

For comparison, results from SPRI 5-fold cross-validations are included in **Figure 3A-B**. However, we note that neither SPRI nor the other methods, PROVEAN (39), PolyPhen-2 (12), PMUT (40), LIST (41), FATHMM (42), EVmutation (13), considered for comparative purposes in the same study (11), adopted a protein-stratified protocol for cross-validations. Instead, a simple protocol that excludes a particular variant from the two sets, but not the other variants of the same protein, was adopted. **Table S2** lists the results from this type of ‘simple’ 5-fold cross-validation for these seven methods and *Rhapsody-2* (and its variants). The table demonstrates that (i) such simple cross-validation overestimates the AUROC and MCC results, and (ii) *Rhapsody-2* outperforms all other methods if the same ‘simple’ cross-validation protocol is adopted for comparative purposes.

Next, we benchmarked *Rhapsody-2* using three independent test sets: AlphaMissense ClinVar, Deciphering Developmental Disorders (DDD) and Cancer Hotspot (19). In this case, we trained *Rhapsody-2* and its variants on subsets of *Rhapsody-2* DB that excluded all proteins whose SAVs were present in the test sets. This resulted in training datasets of 65,382, 107,955, and 110,085 SAVs. The results are presented in **Figure 3C-E**. For comparison, AlphaMissense results are displayed by the *gray bars* in the same plots. However, similarly to SPRI results in panels **A-B**, AlphaMissense results refer to simple tests (in the absence of protein-stratification). Such tests that retain the (other variants) of the tested protein variant in the training set tend to overestimate the performance, as confirmed by application to *Rhapsody-2* and its variants (see **Figure S2)**. Notably, when all methods are equally biased *Rhapsody-2* AUROC values are lower than those of AlphaMissense by 1-2 percentiles, while they surpass those of the next best methods-EVE (9), PrimateAI (48), and VARITY (49) and all other methods used for benchmarking against Missense databases, i.e., Eigen (43), CADDd (44), PolyPhen-2_HVAR (12), ESM1b (45), SIFT (46), PolyPhen-2_HDIV (12), ESM1v (47), VARITY_R_LOO (49), gMVP (50), and REVEL (8). Finally, we note that all these methods, except AlphaMissense, have been trained on limited data available at the time they were developed. Therefore, their relatively lower performance may be due to their training with more limited data.

### Features contributing maximally to classification provide insights into the rationale for SAV predictions by *Rhapsody-2*

The relative contributions of individual features to *Rhapsody-2 (SI* **Table S1**) shed light on the biophysical origins of the predictions. A closer look reveals the features that play a dominant role in determining the effect of the mutation. Notably, among the 100 features, the difference in Position-Specific Independent Counts score (ΔPSIC) (51) between the WT and mutant residue makes the largest contribution (10.30%) to classification. The distributions of ΔPSIC values corresponding to neutral and pathogenic mutations (respective *blue* and *red* histograms in **Figure 4A**) yield medians of 1.71 and 5.28. PSIC reflects the evolutionary conservation at a given position. Its large contribution to *Rhapsody-2* predictions is consistent with the significant effect of sequence conservation or evolution. The second most important feature among sequence evolution-related features is the *z*-score of the Shannon Entropy of the WT residue at the substitution site. This feature contributes 6.54% to classification (**Figure 4B**). It measures the similarity between the WT residue at the substitution site and those of its sequential homologs. A median of -1.01 (highly dissimilar) versus 0.58 (similar) distinguishes the pathogenic and neutral variants. Sequence conservation is closely followed by size-based conservation with 5.3% contribution (see *SI* **Table S1**).

**Figure 4.**
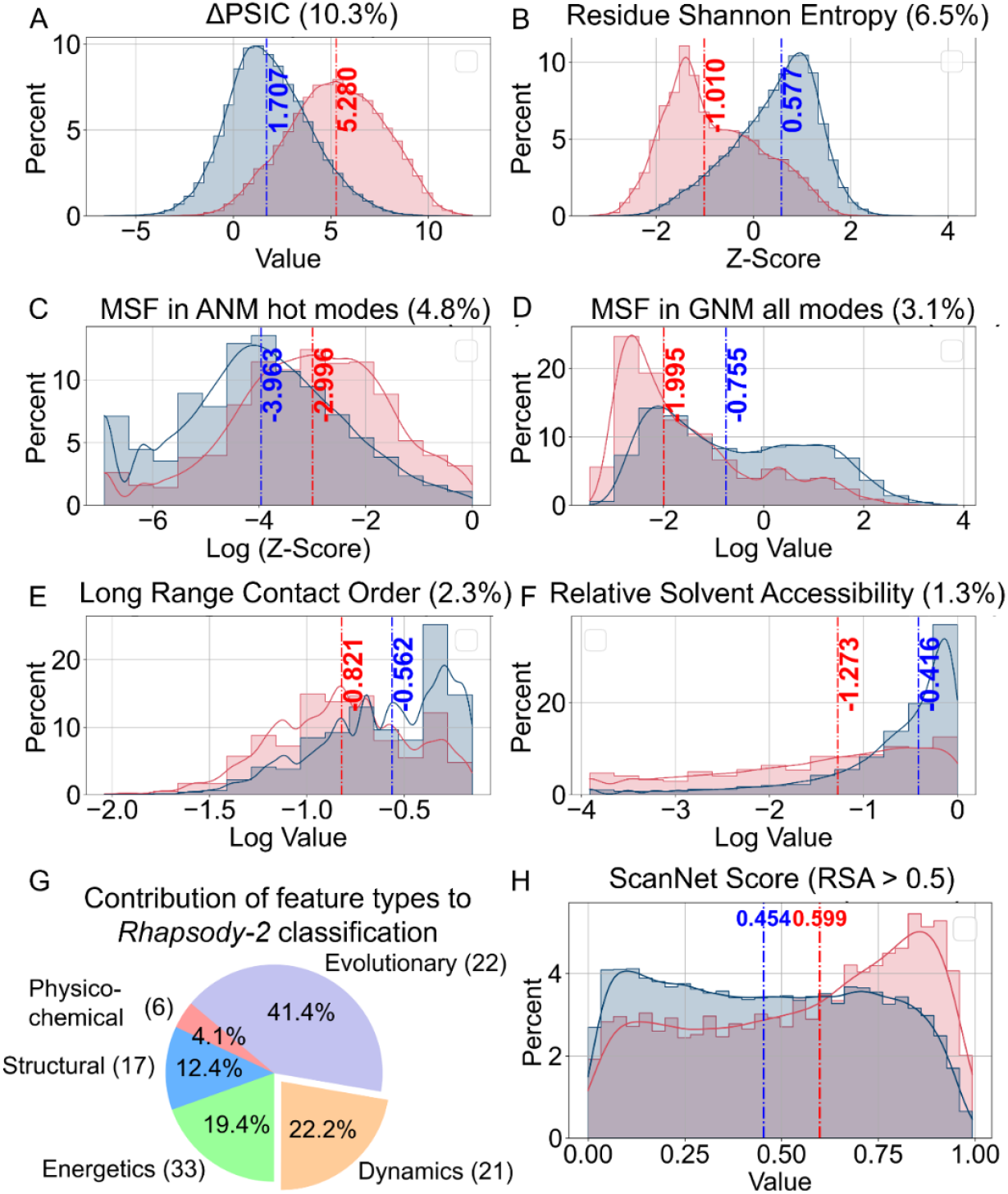
Distribution of selected features distinguished by high discriminatory power between neutral and pathogenic SAVs, and contributions of five categories of features *to Rhapsody-2* predictions. **(A-F)** Histograms of top contributing features in the categories of **(A-B)** sequence evolution, **(C-D)** intrinsic dynamics, and **(E-F)** local interactions. Histograms are presented for neutral (*blue*) and deleterious (*red*) mutations. Median values are indicated by *dashed lines*. **(G)** Percent contributions of different categories of features to *Rhapsody-2* protein-stratified 10-fold classification (**Table S1**). **(H)** Distributions of ScanNet values (probabilities of lying at intermolecular interfaces) for *Rhapsody-2* DB subset of SAVs (48,909 of them) that are solvent exposed (RSA > 0.50).

Among the features describing the protein dynamics, two stand out: participation in high frequency modes of motions, as predicted by the anisotropic network model (ANM) (34), and mean-square-fluctuations (MSFs) under equilibrium conditions, predicted by the Gaussian Network Model (GNM) (33) (**Figure 4C-D**). The *z*-score associated with the former contributes 4.84% to classification. High-frequency modes are manifested by localized fluctuations at the most tightly packed (core) regions, known as kinetically hot sites; these often act as folding nuclei and tend to underlie stability (52-54), hence their strong resistance to substitutions (55). The GNM MSF at the mutation site, on the other hand, contributes 3.13% to classification, ranking second among dynamics features. Its higher median for benign variants, compared to that for pathogenic, shows that mutations at sites that are highly mobile (large MSFs) are more likely to accommodate substitutions, and *vice versa*.

**Figure S3** shows that residues that tend to act as effectors of allosteric signals are more likely to give rise to pathogenicity if mutated. Both GNM- and ANM-based predictions corroborate this behavior. This effect is particularly strong when focusing on GNM soft modes (*upper left panel*). Note that soft (lowest frequency) modes of motion usually entail highly cooperative rearrangements embodying most, if not all, of the structure. It is conceivable that mutations at residues acting as effectors of allosteric signals in the most cooperative modes are prone to cause pathogenicity. Likewise, the residues distinguished by high spatial cross-correlations (positive or negative) with other residues (*bottom right*) show high tendency to give rise to pathogenicity, if mutated.

As to the 17 features associated with local interactions, the most influential is the long-range contact order (LRCO) and relative solvent accessibility (RSA) of the WT residue (**Figure 4E-F**). LRCO quantifies the extent of long-range inter-residue contacts. Contacts are defined as those between residues separated by at least three intervening amino acids along the sequence with farthest atom-atom distance of 8 Å. The LRCO values are normalized with respect to the total number of possible contacts within this distance range. Pathogenic variants exhibit a median LRCO of 0.40, indicating a higher propensity of long-range interactions compared to benign variants (whose median LRCO is 0.18), i.e., amino acids engaged in a higher number of long-range contacts are more susceptible to pathogenicity due the disruption of their extensive interactions. Finally, mutations at buried sites, characterized by low RSA, often disrupt core interactions and structural stability, making them less tolerant to mutations (**Figure 4F**). **Table S3** lists the median values of each feature for neutral and pathogenic SAVs, along with their variance and statistical significance.

The pie chart in **Figure 4G** highlights the relative contributions of different categories of descriptors to *Rhapsody-2* XGBoost classifier, under protein-stratified cross-validation. The evolutionary features make a dominant contribution of 41.4%, followed by dynamics (22.2%) and energetics (19.4%). Note that the model *Rhapsody-2(red)* excluded the energetics, physicochemical, and structural descriptors (except RSA) with minimal loss of predictive power. Therefore, the contribution of different descriptors is model-specific.

Finally, ScanNet (56), the sole feature representing intermolecular interactions, contributes ∼0.5% to classification. ScanNet was used to evaluate the probability of lying at an interface for the examined sites of mutations. **Figure 4H** displays the histograms of ScanNet scores corresponding to pathogenic (*blue*) and benign (*red*) SAVs, using as reference sufficiently exposed (RSA > 0.5) mutation sites. Pathogenic variants have a median score of 0.60, while benign variants score 0.45 (**Figure 4H**), indicating that solvent-exposed residues that make interfacial contacts tend to cause pathogenicity when mutated.

### Influence of the accuracy of AlphaFold2 structural models on the performance of *Rhapsody-2*

For each predicted structure AlphaFold provides a confidence metric, termed “predicted local distance difference test” (pLDDT) score. This metric is assigned per residue and ranges from 0 to 100, with the lower and upper limits referring to the lowest- and highest-confidence predictions. **Figure 5A** shows the distributions of pLDDT scores for pathogenic, neutral and all variants. Notably, neutral SAVs are more common than pathogenic SAVs among proteins predicted with low confidence. In other words, proteins predicted with low confidence show a higher tendency to have neutral SAVs, rather than pathogenic SAVs. This observation could be attributed to the fact that low-confidence regions are usually disordered segments, which, in turn, tend to tolerate mutations, while structured regions are often more tightly packed and less tolerant of mutations. **Figure 5B** shows that the median pLDDT value for neutral residues is 69.04, whereas for pathogenic residues the median is 92.21.

**Figure 5:**
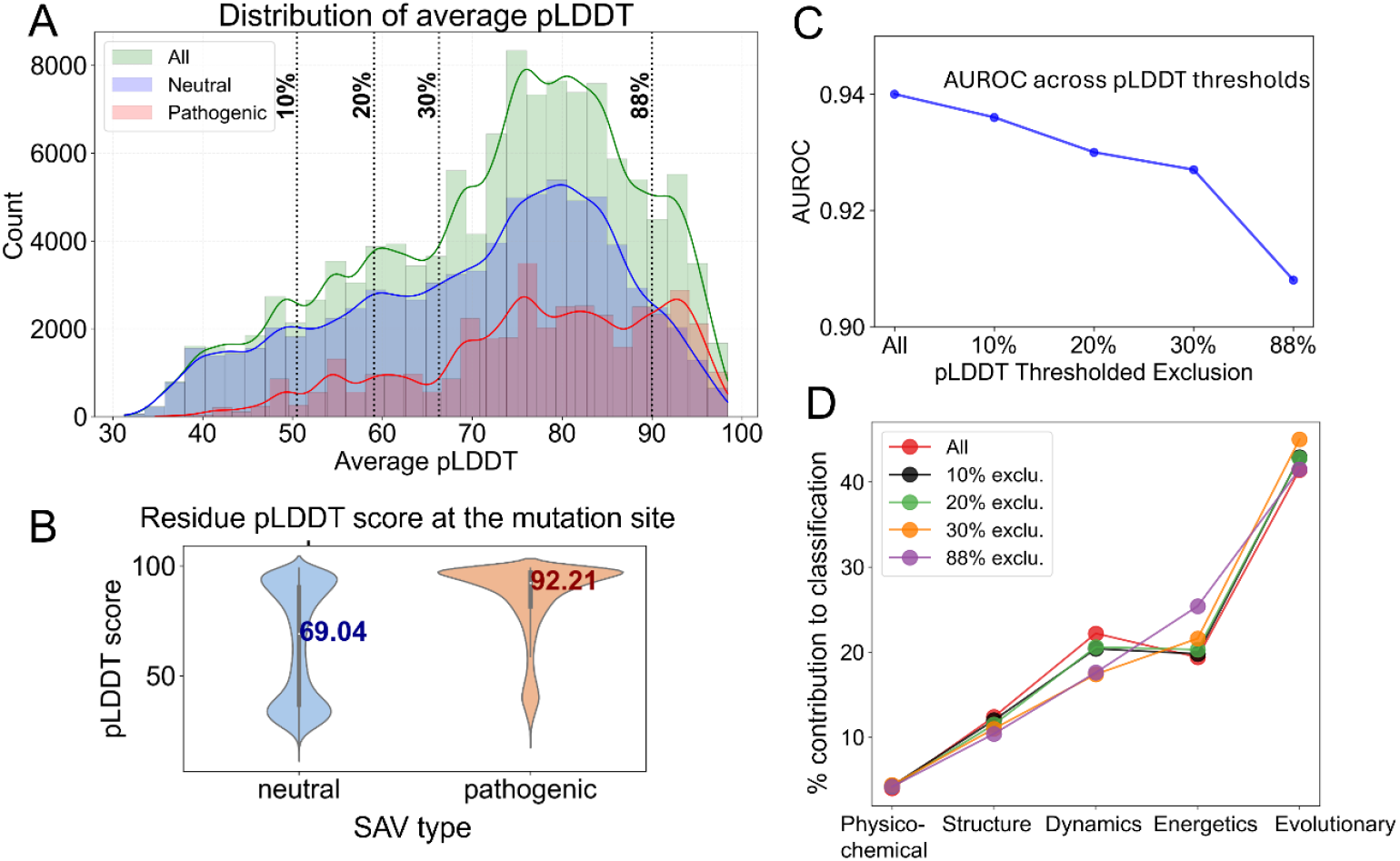
Dependency of *Rhapsody-2* performance on the structural quality of AlphaFold models and size of database used for training. **(A-B)** Distribution of average pLDDT scores for AlphaFold structural models in all (*green*), neutral (*blue*), and pathogenic (*red*) SAVs. The average pLDDT score represents the confidence level of the AlphaFold model averaged over all residues. Kernel density estimates are overlaid on histograms. *Vertical dashed lines* indicate the percentages (10%, 20%, 30%, and 88%) of the *Rhapsody-2* DB that are excluded from the training dataset, corresponding to respective pLDDT lowest thresholds of 50.5, 59.1, 66.3, and 90.0. **(B)** Distribution of pLDDT scores at mutation sites, shown by violin plots for the subsets of neutral and pathogenic SAVs. (**C**) AUROC values as a function of pLDDT threshold. **(D)** Contribution of different categories of descriptors to classification performance at different pLDDT thresholds.

To better understand how the quality of AlphaFold DB-extracted structural data influences prediction performance, we repeated the computations by excluding from the *Rhapsody-2* DB those SAVs which correspond to low-confidence structural models, mainly those with the lowest 10%, 20% and 30% pLDDT scores. We further examined how *Rhapsody-2* performed upon inclusion of structural models with pLDDT scores ≥ 90 or higher, which required the elimination of 88% of the SAVs in the training dataset. As shown in **Figure 5C**, 10-fold protein-stratified cross-validations performed with increasingly smaller sets of structural models decreased the AUROC from 0.940 (original *Rhapsody-2* with all structures) to 0.908 (for the restricted dataset of structures, pLDDT score ≥ 90). This decline could be attributed to the significant decrease in the size of the training dataset (from 117,525 to 14,270 SAVs).

The contribution of specific groups of features to pathogenicity predictions were found to show minor dependencies on the subsets of structures included in the training datasets (**Figure 5D)**. Dynamics features exhibited the highest sensitivity; the removal of low-confidence structural models reduced the percent contribution of dynamic features from 22.2% to 17.6%). This seemingly counter-intuitive trend could be attributed to the exclusion of disordered or highly flexible SAVs modeled as extended loops, which would be otherwise classified as neutral by dynamics features. Most of these SAVs are indeed experimentally detected to be neutral (see **Fig 5A**). Thus, their exclusion reduces the overall AUROC values, as well as the contribution of dynamics features to classification. Energetics contributions, on the other hand, increased from 19.2% to 25.4%, suggesting that improved structural precision allowed for relatively more accurate quantification of interactions.

### A case study illustrating the interpretability of Rhapsody-2 predictions

To illustrate how the outputs *from Rhapsody-2* can help make inferences on the origin of pathogenicity, we present a case study, mainly four pathogenic variants reported for phosphatidylinositol 4,5-bisphosphate 3-kinase catalytic subunit a (PIK3CA) (**Figure 6A**) in the AlphaMissense ClinVar dataset. The α-subunit of PIK3s assumes different conformations in inactive and active forms, which are essential to its catalytic activity (56). The C-terminal end of the catalytic domain of PIK3s “shields” the ATP binding site and is thought to play a role in regulating catalysis, by undergoing a significant conformational change (57). The *in silico* saturation mutagenesis map generated for PIK3CA (selected regions) using *Rhapsody-2* is presented in *SI* **Figure S4**.

**Figure 6.**
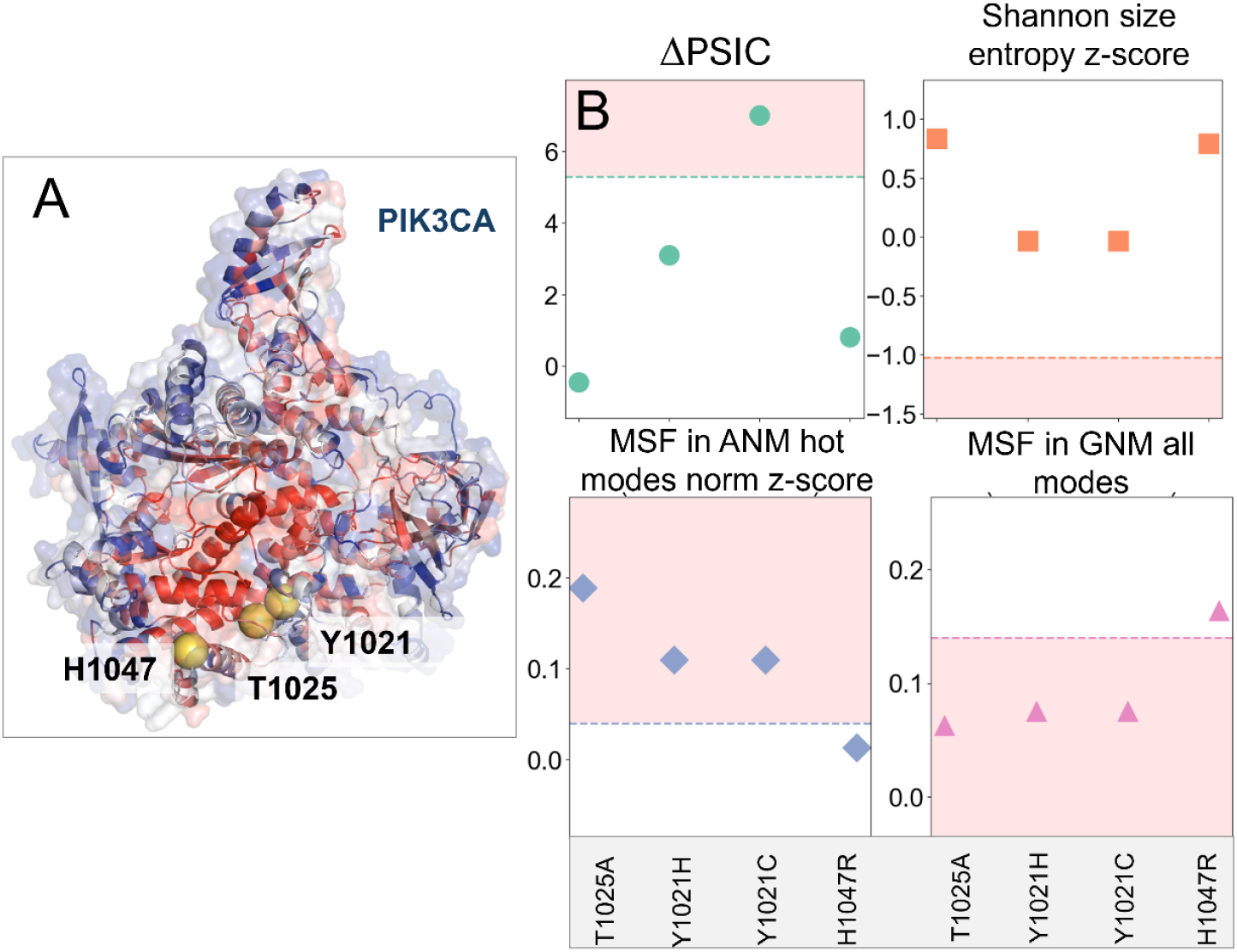
Evaluation of the origin of pathogenicity based on the SAV features examined with respect to median values for pathogenic behavior, illustrated for PIK3CA four mutations. **(A)** Ribbon diagram of PIK3CA color-coded by residue-averaged pathogenicity profile as predicted by *in silico* saturation mutagenesis map (**Figure S4**) generated by *Rhapsody-2* (*red*). The (average) pathogenicity values range from 0 (neutral, in *blue*) to 1(pathogenic, in *red*). The mutated residues are shown in *yellow spheres*. **(B)** Key evolutionary and dynamics features (ordinate) evaluated for the four mutations (abscissa) reported to be pathogenic in ClinVar. Median values of these features (for all DB SAVs) delimit the regions shaded in *pink*, that are strongly pathogenic. Intrinsic dynamics (*bottom two plots*) play a significant role in explaining the pathogenicity of three mutations.

*Rhapsody-2*(*red*) accurately predicts all four mutations T1025A, Y1021H, Y1021C, and H1047R to be pathogenic. **Figure 6B** displays the values of four dominant features (ΔPSIC, size entropy *z*-score, MSF in ANM high frequency modes and MSF in all GNM modes) for these mutations, compared to the median values for all pathogenic SAVs (**Table S3**). The regions shaded in *pink* indicate the pathogenic regions, based on the medians used as threshold. ΔPSIC values for three out of four mutants are below the median of 5.28 (**Figure 6A**), indicating that ΔPSIC was influential only for one of the mutations (Y1021C) and not for the other three. Likewise, the size-based Shannon entropies of the four mutants do not lie within the pathogenic region, indicating that the size change was *not* a factor underlying the observed (and predicted) pathogenicity. Notably, the MSFs in both the ANM hot modes and the GNM all-modes show that T1025A, Y1021H, and Y1021C lie in the pathogenic regime, pointing to the role on intrinsic dynamics in pathogenicity prediction. As to the fourth mutation, H1047R, other features appear to have determined the fate of this mutation, while proximity to the pathogenicity median of the two dynamics features may have somewhat contributed to the classification.

## Discussion and Conclusion

The goal of the present study was 2-fold: to deliver a comprehensible and transparent tool for accurately predicting the pathogenicity of SAVs using not only ‘statistical’ data on sequence patterns but also biophysical data on structure, dynamics and interactions; and to provide a platform for mechanistic interpretation of the predictions based on these biophysical features. *Rhapsody-2* meets both goals. It is a comprehensible tool that provides insights into the biophysical basis of disease-causing mutations. Its ability to integrate and analyze diverse features helps gain a deeper understanding of the molecular basis of dysfunction and develop more effective therapies. A few noteworthy observations are in order.

*High performance of reduced models based different categories of descriptors*. Our computations showed that reduced models based on ∼20 descriptors in a given category (sequence evolution, structure, *or* dynamics) could achieve reasonable performances, despite being trained on different features. This observation points to the interdependence of the descriptors in different categories, consistent with the fundamental tenet: sequence determines structure, which, in turn, defines the equilibrium dynamics or the collective motions (predicted here by the ENMs). Our earlier large-scale analysis (38) indeed showed how sequence evolution goes hand in hand with structural dynamics: e.g., evolutionarily conserved regions exhibit minimal displacements in the global modes and peaks in local modes (52); whereas sequentially variable regions are structurally variable too. Among dynamics descriptors, involvement of a residue in the highest frequency modes turn out to be the strongest determinant of pathogenicity (**Table S1**), consistent with the high tendency of these residues to be evolutionarily conserved.

Yet, among these reduced models, *Rhapsody-2(evo)* outperformed others. This is reminiscent of the success of other sequence-based models such as EVE (9) and EVmutation (13). Dynamics-based descriptors were the 2^nd^ highest contributor (22.2%) to *Rhapsody-2* classification, after sequence evolution (41.4%) (**Figure 4G**). Not surprisingly, a reduced model, *Rhapsody-2(red)*, trained on these two categories of descriptors (plus solvent accessibility; a total of 44 features) was distinguished as a powerful predictor **(Figures 2** and **3**). Its performance was comparable to that of *Rhapsody-2* (trained on 100 descriptors), especially when benchmarked against specialized databases (**Figure 3**). The fact that *Rhapsody-2(red)* outperformed both *Rhapsody-2(evo)* and *Rhapsody-2(dyn)* indicates that these descriptors, despite their interdependencies, do contain complementary information. Furthermore, there is a merit to consider both sets of predictors as illustrated in **Figure 6**.

*Rhapsody-2* permits us to assess the molecular bases of the predictions. To this aim, one may refer to the histograms of dominant features generated for neutral and pathogenic SAVs, or the tabulated median values of all features for neutral and pathogenic mutations (**Table S3**) and examine how the individual SAV’s features compare to those data, as illustrated in **Figure 6**.

### Influence of structural models extracted from the AlphaFold DB

The superior performance of *Rhapsody-2* (and AlphaMissense) compared to other pathogenicity predictors can be largely attributed to the vast amount of structural data made available through AlphaFold DB. ESM1b and AlphaMissense brought a paradigm shift by leveraging advanced sequence and structure data, which earlier methods did not benefit from. The other methods could have shown comparable success if they had access to the same data. Inclusion of a larger number of AlphaFold DB structural models, even when these were low confidence models, helped increase the AUROC. This can be seen from the decrease in AUROC values (**Figure 5C**) as increasingly larger subsets of structures with relatively low confidence levels were excluded from the training dataset. In the extreme case of a dataset composed of only highly accurate (pLDDT > 90) structural models, the AUROC from protein-stratified 10-fold cross-validations dropped to 0.908 (as opposed to 0.940 for the complete set). Thus, the benefit of learning from a large dataset more than offsets the possible detrimental effects of poorly defined structural regions. Notably, most of the low-confidence structures have neutral mutations based on current experiments (**Figure 5A**), and because they are modeled by AlphaFold as extended or disordered loops they are also classified as neutral by dynamics descriptors.

### Future challenges

Moving forward, there are still areas that demand further work, especially in the case of specialized datasets. For example, when benchmarked against *AlphaMissense* DDD, all methods showed an AUROC < 0.81. It may be necessary to develop more system-specific classifiers, such as those customized to membrane proteins, or multimeric proteins, or manifested in different cell/tissue contexts. *Rhapsody-2* and most pathogenicity predictors are based on the properties of the SAVs themselves without explicit consideration of systems/context-dependent effects. Yet, the interactions in the cellular environment and how the mutations impact those interactions may be a serious consideration, and there are recent studies that focus on the impact of mutations on protein-protein interaction interfaces (57, 58).

Other challenges include the characterization of the effects of multiple mutations, those of insertion/deletions, or association/dissociation of entire domains/subunits. Intrinsically disordered protein (IDP) segments remain as challenging tasks that await further work. In addition, post-translational modifications, transmembrane topology and small-molecule, metal or ion binding, are crucial to function (15, 16, 59), which may need attention in future pathogenicity predictors. We have not explicitly analyzed intrinsically disordered protein (IDP) regions while training or testing *Rhapsody-2*, even though the pLDDT score distributions suggest that *Rhapsody-2* DB contains SAVs from IDP regions. Given the functional significance of IDPs in cellular context (60), a critical assessment of pathogenicity predictions for SAVs at IDP regions might be an important future direction.

Overall, *Rhapsody-2* is a comprehensible tool that not only distinguishes pathogenic and neutral mutations but also provides insights into the biophysical basis of disease-causing mutations. Its ability to integrate and analyze diverse data is essential to gaining a deeper understanding of the impact of mutations and providing guidance toward developing more effective therapies.

## Materials and Methods

Detailed descriptions of the datasets, features, XGBoost classifier specifications, and performance metrics used in this study are presented in the *SI*-Appendix. The datasets and codes are available at https://github.com/anupam-banerjee/rhapsody-2.

## Acknowledgment

Support from NIH award R01GM139297 is gratefully acknowledged by IB.

## References

1. S. Stefl, H. Nishi, M. Petukh, A. R. Panchenko, E. Alexov, Molecular mechanisms of disease-causing missense mutations. J Mol Biol 425, 3919–3936 (2013).

2. M. J. Landrum et al., ClinVar: improving access to variant interpretations and supporting evidence. Nucleic Acids Res 46, D1062–d1067 (2018).

3. A. Kamburov et al., Comprehensive assessment of cancer missense mutation clustering in protein structures. Proc Natl Acad Sci U S A 112, E5486–5495 (2015).

4. M. H. Meisler, J. A. Kearney, Sodium channel mutations in epilepsy and other neurological disorders. J Clin Invest 115, 2010–2017 (2005).

5. H. Zhao et al., SIGMA leverages protein structural information to predict the pathogenicity of missense variants. Cell Rep Methods 4, 100687 (2024).

6. S. Ittisoponpisan et al., Can Predicted Protein 3D Structures Provide Reliable Insights into whether Missense Variants Are Disease Associated? J Mol Biol 431, 2197–2212 (2019).

7. R. Vaser, S. Adusumalli, S. N. Leng, M. Sikic, P. C. Ng, SIFT missense predictions for genomes. Nat Protoc 11, 1–9 (2016).

8. N. M. Ioannidis et al., REVEL: An Ensemble Method for Predicting the Pathogenicity of Rare Missense Variants. Am J Hum Genet 99, 877–885 (2016).

9. J. Frazer et al., Disease variant prediction with deep generative models of evolutionary data. Nature 599, 91–95 (2021).

10. J. Lai, J. Yang, E. D. Gamsiz Uzun, B. M. Rubenstein, I. N. Sarkar, LYRUS: a machine learning model for predicting the pathogenicity of missense variants. Bioinform Adv 2, vbab045 (2022).

11. B. Wang et al., Structure-based pathogenicity relationship identifier for predicting effects of single missense variants and discovery of higher-order cancer susceptibility clusters of mutations. Brief Bioinform 24 (2023).

12. I. Adzhubei, D. M. Jordan, S. R. Sunyaev, Predicting functional effect of human missense mutations using PolyPhen-2. Curr Protoc Hum Genet Chapter 7, Unit7.20 (2013).

13. T. A. Hopf et al., Mutation effects predicted from sequence co-variation. Nat Biotechnol 35, 128–135 (2017).

14. E. Capriotti et al., WS-SNPs&GO: a web server for predicting the deleterious effect of human protein variants using functional annotation. BMC Genomics 14 Suppl 3, S6 (2013).

15. B. Li et al., Automated inference of molecular mechanisms of disease from amino acid substitutions. Bioinformatics 25, 2744–2750 (2009).

16. V. Pejaver et al., Inferring the molecular and phenotypic impact of amino acid variants with MutPred2. Nat Commun 11, 5918 (2020).

17. H. Qi et al., MVP predicts the pathogenicity of missense variants by deep learning. Nat Commun 12, 510 (2021).

18. N. Brandes, G. Goldman, C. H. Wang, C. J. Ye, V. Ntranos, Genome-wide prediction of disease variant effects with a deep protein language model. Nat Genet 55, 1512–1522 (2023).

19. J. Cheng et al., Accurate proteome-wide missense variant effect prediction with AlphaMissense. Science 381, eadg7492 (2023).

20. J. Jumper et al., Highly accurate protein structure prediction with AlphaFold. Nature 596, 583–589 (2021).

21. A. Banerjee, S. Saha, N. C. Tvedt, L. W. Yang, I. Bahar, Mutually beneficial confluence of structure-based modeling of protein dynamics and machine learning methods. Curr Opin Struct Biol 78, 102517 (2023).

22. L. Orellana et al., Oncogenic mutations at the EGFR ectodomain structurally converge to remove a steric hindrance on a kinase-coupled cryptic epitope. Proceedings of the National Academy of Sciences 116, 10009–10018 (2019).

23. L. Ponzoni, I. Bahar, Structural dynamics is a determinant of the functional significance of missense variants. Proc Natl Acad Sci U S A 115, 4164–4169 (2018).

24. L. Ponzoni, D. A. Penaherrera, Z. N. Oltvai, I. Bahar, Rhapsody: predicting the pathogenicity of human missense variants. Bioinformatics 36, 3084–3092 (2020).

25. I. A. Adzhubei et al., A method and server for predicting damaging missense mutations. Nat Methods 7, 248–249 (2010).

26. J. Bendl et al., PredictSNP: robust and accurate consensus classifier for prediction of disease-related mutations. PLoS Comput Biol 10, e1003440 (2014).

27. M. X. Li et al., Predicting mendelian disease-causing non-synonymous single nucleotide variants in exome sequencing studies. PLoS Genet 9, e1003143 (2013).

28. A. Mottaz, F. P. David, A. L. Veuthey, Y. L. Yip, Easy retrieval of single amino-acid polymorphisms and phenotype information using SwissVar. Bioinformatics 26, 851–852 (2010).

29. J. Thusberg, A. Olatubosun, M. Vihinen, Performance of mutation pathogenicity prediction methods on missense variants. Hum Mutat 32, 358–368 (2011).

30. D. G. Grimm et al., The evaluation of tools used to predict the impact of missense variants is hindered by two types of circularity. Hum Mutat 36, 513–523 (2015).

31. S. Zhang et al., ProDy 2.0: increased scale and scope after 10 years of protein dynamics modelling with Python. Bioinformatics 37, 3657–3659 (2021).

32. A. Bakan, L. M. Meireles, I. Bahar, ProDy: protein dynamics inferred from theory and experiments. Bioinformatics 27, 1575–1577 (2011).

33. I. Bahar, A. R. Atilgan, B. Erman, Direct evaluation of thermal fluctuations in proteins using a single-parameter harmonic potential. Fold Des 2, 173–181 (1997).

34. A. R. Atilgan et al., Anisotropy of fluctuation dynamics of proteins with an elastic network model. Biophys J 80, 505–515 (2001).

35. E. Eyal, G. Lum, I. Bahar, The anisotropic network model web server at 2015 (ANM 2.0). Bioinformatics 31, 1487–1489 (2015).

36. B. Buchfink, K. Reuter, H. G. Drost, Sensitive protein alignments at tree-of-life scale using DIAMOND. Nat Methods 18, 366–368 (2021).

37. T. Chen, C. Guestrin (2016) Xgboost: A scalable tree boosting system. in Proceedings of the 22nd acm sigkdd international conference on knowledge discovery and data mining, pp 785–794.

38. Y. Liu, I. Bahar, Sequence evolution correlates with structural dynamics. Mol Biol Evol 29, 2253–2263 (2012).

39. Y. Choi, A. P. Chan, PROVEAN web server: a tool to predict the functional effect of amino acid substitutions and indels. Bioinformatics 31, 2745–2747 (2015).

40. V. Lopez-Ferrando, A. Gazzo, X. de la Cruz, M. Orozco, J. L. Gelpi, PMut: a web-based tool for the annotation of pathological variants on proteins, 2017 update. Nucleic Acids Res 45, W222–W228 (2017).

41. N. Malhis, M. Jacobson, S. J. M. Jones, J. Gsponer, LIST-S2: taxonomy based sorting of deleterious missense mutations across species. Nucleic Acids Res 48, W154–W161 (2020).

42. H. A. Shihab et al., Predicting the functional, molecular, and phenotypic consequences of amino acid substitutions using hidden Markov models. Hum Mutat 34, 57–65 (2013).

43. I. Ionita-Laza, K. McCallum, B. Xu, J. D. Buxbaum, A spectral approach integrating functional genomic annotations for coding and noncoding variants. Nat Genet 48, 214–220 (2016).

44. P. Rentzsch, D. Witten, G. M. Cooper, J. Shendure, M. Kircher, CADD: predicting the deleteriousness of variants throughout the human genome. Nucleic Acids Res 47, D886–d894 (2019).

45. A. Rives et al., Biological structure and function emerge from scaling unsupervised learning to 250 million protein sequences. Proc Natl Acad Sci U S A 118 (2021).

46. P. C. Ng, S. Henikoff, SIFT: Predicting amino acid changes that affect protein function. Nucleic Acids Res 31, 3812–3814 (2003).

47. S. Mansoor, M. Baek, D. Juergens, J. L. Watson, D. Baker, Zero-shot mutation effect prediction on protein stability and function using RoseTTAFold. Protein Sci 32, e4780 (2023).

48. L. Sundaram et al., Predicting the clinical impact of human mutation with deep neural networks. Nat Genet 50, 1161–1170 (2018).

49. Y. Wu, R. Li, S. Sun, J. Weile, F. P. Roth, Improved pathogenicity prediction for rare human missense variants. Am J Hum Genet 108, 1891–1906 (2021).

50. H. Zhang, M. S. Xu, X. Fan, W. K. Chung, Y. Shen, Predicting functional effect of missense variants using graph attention neural networks. Nat Mach Intell 4, 1017–1028 (2022).

51. S. R. Sunyaev et al., PSIC: profile extraction from sequence alignments with position-specific counts of independent observations. Protein engineering 12, 387–394 (1999).

52. I. Bahar, A. R. Atilgan, M. C. Demirel, B. Erman, Vibrational Dynamics of Folded Proteins: Significance of Slow and Fast Motions in Relation to Function and Stability. Physical Review Letters 80, 2733–2736 (1998).

53. M. C. Demirel, A. R. Atilgan, R. L. Jernigan, B. Erman, I. Bahar, Identification of kinetically hot residues in proteins. Protein Sci 7, 2522–2532 (1998).

54. A. Rader, I. Bahar, Folding core predictions from network models of proteins. Polymer 45, 659–668 (2004).

55. A. Banerjee, I. Bahar, Structural Dynamics Predominantly Determine the Adaptability of Proteins to Amino Acid Deletions. Int J Mol Sci 24 (2023).

56. J. Tubiana, D. Schneidman-Duhovny, H. J. Wolfson, ScanNet: an interpretable geometric deep learning model for structure-based protein binding site prediction. Nat Methods 19, 730–739 (2022).

57. G. Hanna, T. Khanna, S. A. Islam, A. David, M. J. E. Sternberg, Missense3D-TM: Predicting the Effect of Missense Variants in Helical Transmembrane Protein Regions Using 3D Protein Structures. J Mol Biol 436, 168374 (2024).

58. C. Pennica, G. Hanna, S. A. Islam, M. J. E. Sternberg, A. David, Missense3D-PPI: A Web Resource to Predict the Impact of Missense Variants at Protein Interfaces Using 3D Structural Data. J Mol Biol 435, 168060 (2023).

59. V. Pejaver, S. D. Mooney, P. Radivojac, Missense variant pathogenicity predictors generalize well across a range of function-specific prediction challenges. Hum Mutat 38, 1092–1108 (2017).

60. A. S. Holehouse, B. B. Kragelund, The molecular basis for cellular function of intrinsically disordered protein regions. Nat Rev Mol Cell Biol 25, 187–211 (2024).

