## Supplementary Information for "Accurate Identification and Mechanistic Evaluation of Pathogenic Missense Variants with *Rhapsody-2*"

*for*

**This PDF file includes:**

Supplementary Methods
Tables S1 to S3
Figures S1 to S4
Supplementary References

### SUPPLEMENTARY METHODS

***Rhapsody-2* Database (DB) curation and variant classification.** We used the ClinVar database (1) as of December 2023 to develop the *Rhapsody-2* DB. This database encompasses 117,525 SAVs across 12,094 human proteins. Of these variants, 34% were classified as pathogenic. Variants labeled with ClinVar significance (CLNSIG) as “Benign” or “Likely benign” were categorized as benign (negative), and those labeled as “Pathogenic” or “Likely pathogenic” were categorized as pathogenic (positive). Entries marked with “Conflicting classifications of pathogenicity” were excluded from this study. For each variant, the corresponding protein structure in the AlphaFold database (2) was utilized to compute various features

**Clinical Benchmark Databases**. The UnifyPDBFull and UnifyPDBAcceptable datasets were introduced in 2023 by Want et al., while developing the structure-based pathogenicity relationship identifier (SPRI) (3). The former contains 4,231 deleterious variants and 2,791 neutral variants derived from 252 single-chain proteins with full structural coverage; and the latter 5,999 deleterious and 3485 neutral variants, derived from 377 proteins with 444 polypeptide chains with full and partial structural coverage. UnifyPDBFull is a subset of the UnifyPDBAcceptable dataset.

AlphaMissense test sets: The AlphaMissense (4) ClinVar test set comprises 9,462 pathogenic and 9,462 benign variants sourced from 999 proteins, ensuring a balanced representation of positive and negative variants per gene within the ClinVar database. The AlphaMissense Deciphering Developmental Disorders (DDD) benchmark database (5) includes *de novo* variants identified in patients from the DDD cohort, along with variants from healthy controls. It incorporates 353 patient variants and 57 control variants derived from 215 genes relevant to developmental disorders as studied by the DDD project. The AlphaMissense Cancer Hotspot database comprises of 868 missense variants in hotspots from 202 cancer driver genes (6) versus 1,734 randomly selected negative variants from the DiscovEHR data (7).

**Features for biophysical characterization of missense variants**. As described in the main text, we considered five categories of features, comprising a total of 100 features. The features belonging to each category are detailed below.

*Residue-specific attributes*. We evaluated four residue-specific features for both the wild-type (WT) and mutated amino acids: molecular weight, hydrophobicity score after the Kyte and Doolittle scale (8), secondary structure propensity (9), and group belongingness. The latter classifies the amino acid into hydrophobic, polar, acidic, basic, or aromatic groups. This resulted in a total of eight residue-specific features for each SAV.

*Local interactions*. The close environment of a residue undergoing a point mutation and its interactions with close spatial neighbors in the 3D structure are captured through this group of features. First, the side chain rotameric state at the mutation site is optimized using SCWRL (10) for both the WT and mutant protein. This optimization accommodates the mutated residue and ensures uniformity while refining poor-quality protein structures, if any. The features considered to describe local interactions include: weighted contact number (WCN), weighted hydrophobic contact number (WHbCN), aromatic cluster score (ACS), hydrophobic core score (HCS), hydrophobic buried core score (HBCS), and long-range contact order (LRCO) (11). WCN reflects the density of neighboring residues for the native amino acid. It is the inverse sum of the square of the distance from the WT residue to all other residues based on the geometric center of the residues. It reveals how deeply seated the amino acid is within the protein structure. WHbCN extends WCN by encoding the hydrophobic density of the neighboring residues. ACS measures the maximum number of aromatic residues whose geometric centers lie within 8 Å from each other and from the residue in question. HCS represents the ratio of hydrophobic residues to all residues whose geometric centers lie within 6 Å from the geometric center of the mutated residue. HBCS considers the ratio of buried hydrophobic residues to all residues in the same geometric vicinity. LRCO encodes the long-range interactions of the residue undergoing a mutation.

We examine the ACS, HCS, and HBCS measures for both the native and mutated residues, while WCN, WHbCN, LRCO, and relative solvent accessibility (RSA) (using STRIDE (12)) were considered only for the native amino acid. Additionally, information on salt bridges, disulfide linkages, and hydrogen bonds (calculated using HBPLUS (13)) for both the native and mutated amino acids) is included.

Furthermore, ScanNet (14), an interpretable geometric deep learning model for structure-based protein binding site prediction, was used to identify the likelihood of the mutated residue to be part of an protein-protein interfacial site. The probability value as predicted by ScanNet ranges from 0 to 1, where upper limit corresponds to the highest (100%) probability of lying at an intermolecular interface for the examined residue, and the lower limit means zero probability. In total, 18 features representative of local interactions or environmental effects were considered.

*Intrinsic dynamics*. We computed intrinsic dynamics-based features for the WT protein using elastic network models via the *ProDy* API (15). A total of 21 dynamics-based features were used in the *Rhapsody-2* model and its variants.

First, we employed the Gaussian Network Model (GNM) (16) and Anisotropic Network Model (ANM) (17) to calculate the mean-square fluctuations (MSF) of residues across different sets of modes: all modes, the first 2% of modes (hot), the five lowest-frequency modes (top 5), and the softest modes. Descriptors of these modes included corresponding MSFs, Z-Score MSF, and normalized Z-Score MSF.

Second, perturbation response scanning (PRS) (18, 19) was performed to evaluate the effectiveness and sensitivity of the mutation site for allosteric signal transmission and sensing. PRS features included metrics such as effectiveness, normalized effectiveness, and their Z-Scores, computed for selected GNM and ANM modes.

Third, we analyzed the mean cross-correlation between the mutated residue and all other residues, derived from the GNM cross-correlation matrix and focused on the softest modes.

These features, listed in **Table S1**, were selected for their ability to capture essential aspects of protein dynamics while maintaining computational efficiency and relevance to the functional consequences of mutations.

Finally, we assessed whether the mutated residue acted as a hinge residue using the *calcHinges* function of *ProDy*, identifying residues closest to the cross-over points in the eigenvectors of the GNM global modes. We evaluated this for the first 2% and the first 5 modes to determine the hinge site association of the mutated residue. Overall, 81 dynamics-based attributes were used for constructing the *Rhapsody-2* model and its variants.

*Energetics of the folded state* . We utilized the empirical energy function FoldX (20) to compute the ΔG of folding of both the WT and mutant structures. Additionally, we considered 15 individual components of the FoldX energy function for both the native and mutant structures, resulting in a total of 32 features derived from FoldX. For a detailed list of these components, refer to Table S1. Moreover, we incorporated the predicted change in Gibbs free energy of folding due to the mutant substitution, as provided by PROTSPOM (11).

*Evolutionary features*. These were computed using a multi-step process leveraging existing bioinformatics tools and databases. First, the DIAMOND package (21) facilitated BLAST searches for all human proteins cataloged in the AlphaFold2 database, accommodating fragmented entries as necessary. Multiple sequence alignments (MSAs) were constructed across the NCBI non-redundant (NR) database, requiring a minimum 30% coverage with the target sequence. Using the MSAs, Position Specific Independent Counts (PSIC) (22) were calculated for individual residues, as well as for physicochemical and size groups of both wild-type (WT), mutant (Mut), and the difference in PSIC scores (ΔPSIC), totaling nine distinct features. Shannon Entropy, a measure of sequence variability/conservation, was computed for each residue, with corresponding Z-scores computed for both the individual residues and their groups based on physicochemical properties and size. The computations encompassed both gapped and ungapped MSAs, resulting in twelve features for assessing evolutionary conservation and variability across aligned sequences. We additionally considered the residue substitution score of the WT amino acid by the mutated amino acid from the BLOSUM62 matrix (23), overall summing up to 22 evolutionary features.

**XGBoost classification and cross-validation.** To assess the performance of our model, we employed the XGBoost (24) classifier using the XGBOOST Python package along with 10-fold stratified cross-validation and 10-fold protein-stratified cross-validation (using StratifiedGroupKFold from sklearn). This approach ensures that each fold maintains the same class distribution as the entire dataset, providing a robust evaluation of model performance across different subsets of the data. We split the dataset into features and labels, initializing the StratifiedKFold method with 10 splits to maintain class balance within each fold. Additionally, we applied StratifiedGroupKFold with 10 splits to ensure that proteins in each test set are entirely distinct from those in the corresponding training sets, thereby avoiding data leakage. The classifier was configured with a maximum tree depth of 11 and 750 boosting rounds to enhance model complexity while mitigating overfitting, coupled with a conservative learning rate of 0.02. We leveraged GPU acceleration ('cuda') when available to expedite model training. Subsampling strategies included colsample_bytree, colsample_bylevel, and colsample_bynode set to 0.7, 0.8 and 0.7 respectively for *Rhapsody-2* (and 0.7, 0.8 and 0.9 for all other classifiers with feature subsets), optimizing feature selection at various levels of tree construction. Regularization techniques incorporated a gamma value of 5 for *Rhapsody-2* (and 1 for all other classifiers) to control split complexity and min_child_weight set to 1 for stability in leaf node partitioning. Further regularization was applied with reg_alpha = 0.5 (L1) and reg_lambda = 5 (L2) to penalize model complexity and prevent overfitting. Additionally, scale_pos_weight = 2 was adopted to address class imbalance and ensuring robust performance across different datasets used for evaluation. The same parameters were used across all training databases.

**Performance evaluation.** The performance of the XGBOOST classifier was evaluated using various metrics, as summarized below.

*Accuracy*- This metric measures how often the model makes correct predictions across all classes. It is calculated by dividing the sum of true positives (TP) and true negatives (TN) by the total number of instances including false positives (FPs) and false negatives (FNs)

$$Accuracy = \frac{TP\ +\ TN}{TP\ +\ TN\ \_FP\ +\ FN}$$

*Precision*- Precision indicates the accuracy of positive predictions, showcasing the proportion of correctly identified positive instances out of all instances labeled as positive, i.e.

$$Precision = \frac{TP}{TP + FP}$$

*Recall (Sensitivity)*- This metric measures the ability of the model to capture all relevant instances. It is calculated as the ratio of true positives to the sum of true positives and false negatives.

$$Recall = \frac{TP}{TP\ +\ FN}$$

*F1 Score*- The F1 Score represents the harmonic mean of precision and recall. It provides a balanced evaluation where higher values indicate better performance.

$$F1\,Score = \frac{2 \times Precision \times Recall}{Precision + Recall}$$

*Area Under the Receiver Operating Characteristic Curve (AUROC)*- AUROC assesses the model's ability to distinguish between classes by plotting the TP rate against the FP rate. Higher AUROC values indicate better discrimination performance.

$$True\ Positive\ Rate = \frac{TP}{TP\ +\ FN}$$

$$False\ Positive\ Rate = \frac{FP}{FP + TN}$$

*Area Under the Precision-Recall Curve (AUPRC)*- AUPRC evaluates the precision-recall trade-off across different thresholds. This metric is particularly useful for imbalanced datasets, where precision and recall may vary.

*Matthews Correlation Coefficient (MCC)*- MCC is a statistical measure used to evaluate the performance of binary classification models, particularly in situations where classes are imbalanced, defined as

$$MCC = \frac{(TN \times TP) - (FN \times FP)}{\sqrt{(TP+FP)(TP+FN)(TN+FP)(TN+FN)}}.$$

MCC provides a balanced assessment of a model's predictive ability. A MCC score of +1 indicates perfect prediction, 0 indicates no better than random prediction, and -1 indicates total disagreement between prediction and observation. It is widely favored for its ability to reflect a classifier's performance across all classes, making it a robust metric in biomedical and machine learning contexts.

**Evaluation of the importance/contribution of individual features.** The percent contributions of all features used in *Rhapsody-2* were averaged over the 10-fold cross-validation computations to identify which features contributed most significantly to the decision-making process of the model. This comprehensive evaluation framework provided a detailed understanding of the XGBoost classifier's performance in relation to its underlying features.

### SUPPLEMENTARY TABLES

**Table S1.** *Rhapsody-2* features (arranged by type and sorted by contribution to classification in each type) and their contribution to classification

| Feature | Feature Name | Feature type | Contribution (%) | |
|---|---|---|---|---|
| | | | 10-fold stratified | 10-fold protein stratified |
| 1 | WT-Mut Seq_PSIC (with gaps) | Evolutionary | 10.30 | 9.31 |
| 2 | WT zscore Seq_Shannon Entropy (without gaps) | Evolutionary | 6.54 | 6.15 |
| 3 | WT Size Group zcore Seq_Shannon Entropy (without gaps) | Evolutionary | 5.31 | 6.06 |
| 4 | Mut Seq_PSIC (with gaps) | Evolutionary | 3.46 | 2.74 |
| 5 | WT Size Group Seq_Shannon Entropy (without gaps) | Evolutionary | 2.08 | 3.36 |
| 6 | BLOSSUM (WT, Mut) | Evolutionary | 2.00 | 2.01 |
| 7 | WT Size Group Seq_PSIC (with gaps) | Evolutionary | 1.92 | 1.89 |
| 8 | WT PhyChem Group zcore Seq_Shannon Entropy (without gaps) | Evolutionary | 1.55 | 1.70 |
| 9 | WT Seq_PSIC (with gaps) | Evolutionary | 1.34 | 1.17 |
| 10 | WT PhyChem Group Seq_Shannon Entropy (without gaps) | Evolutionary | 1.12 | 0.56 |
| 11 | WT Seq_Shannon Entropy (without gaps) | Evolutionary | 1.11 | 0.93 |
| 12 | WT-Mut Size Group Seq_PSIC (with gaps) | Evolutionary | 0.92 | 0.79 |
| 13 | WT-Mut PhyChem Group Seq_PSIC (with gaps) | Evolutionary | 0.83 | 0.76 |
| 14 | WT PhyChem Group Seq_PSIC (with gaps) | Evolutionary | 0.78 | 0.75 |
| 15 | WT zscore Seq_Shannon Entropy (with gaps) | Evolutionary | 0.43 | 0.42 |
| 16 | WT Seq_Shannon Entropy (with gaps) | Evolutionary | 0.42 | 0.44 |
| 17 | WT Size Group Seq_Shannon Entropy (with gaps) | Evolutionary | 0.42 | 0.43 |
| 18 | WT PhyChem Group Seq_Shannon Entropy (with gaps) | Evolutionary | 0.40 | 0.41 |
| 19 | WT PhyChem Group zcore Seq_Shannon Entropy (with gaps) | Evolutionary | 0.40 | 0.39 |
| 20 | WT Size Group zcore Seq_Shannon Entropy (with gaps) | Evolutionary | 0.40 | 0.41 |
| 21 | Mut PhyChem Group Seq_PSIC (with gaps) | Evolutionary | 0.39 | 0.37 |
| 22 | Mut Size Group Seq_PSIC (with gaps) | Evolutionary | 0.39 | 0.36 |
| **Sum** | | | **42.51** | **41.41** |
| 23 | res_MSF_zscore (ANM hot) | Dynamics | 4.84 | 5.14 |
| 24 | res_MSF (GNM all) | Dynamics | 3.13 | 2.76 |
| 25 | res_MSF (ANM hot) | Dynamics | 2.21 | 2.48 |
| 26 | res_MSF (GNM soft) | Dynamics | 1.58 | 1.72 |
| 27 | res_MSF_norm_zscore (GNM soft) | Dynamics | 0.94 | 0.95 |
| 28 | res_MSF_norm_zscore (ANM hot) | Dynamics | 0.90 | 1.03 |
| 29 | res_MSF_norm_zscore (GNM all) | Dynamics | 0.90 | 0.90 |
| 30 | res_MSF (GNM top5) | Dynamics | 0.76 | 0.79 |

| | | | | |
|---|---|---|---|---|
| 31 | res_effectiveness (GNM soft) | Dynamics | 0.64 | 0.65 |
| 32 | res_norm_zscore_effectiveness (GNM all) | Dynamics | 0.61 | 0.59 |
| 33 | res_MSF_norm_zscore (GNM hot) | Dynamics | 0.58 | 0.60 |
| 34 | res_avg_meancorr (GNM soft) | Dynamics | 0.56 | 0.54 |
| 35 | res_MSF (ANM all) | Dynamics | 0.54 | 0.54 |
| 36 | res_MSF (ANM soft) | Dynamics | 0.51 | 0.50 |
| 37 | res_zscore_effectiveness (GNM all) | Dynamics | 0.50 | 0.50 |
| 38 | res_effectiveness (GNM all) | Dynamics | 0.50 | 0.51 |
| 39 | res_effectiveness (ANM all) | Dynamics | 0.45 | 0.43 |
| 40 | res_norm_zscore_sensitivity (ANM all) | Dynamics | 0.44 | 0.43 |
| 41 | res_MSF_zscore (GNM hot) | Dynamics | 0.42 | 0.41 |
| 42 | res_zscore_sensitivity (ANM top5) | Dynamics | 0.42 | 0.41 |
| 43 | res_hinge_binary (GNM soft) | Dynamics | 0.32 | 0.32 |
| **Sum** | | | **21.75** | **22.20** |
| 44 | WT Disulfide FoldX | Energetics | 0.89 | 0.85 |
| 45 | Mut Disulfide FoldX | Energetics | 0.87 | 0.84 |
| 46 | Mut Entropy MC FoldX | Energetics | 0.78 | 0.82 |
| 47 | WT Entropy MC FoldX | Energetics | 0.75 | 0.76 |
| 48 | WT Torsion FoldX | Energetics | 0.65 | 0.66 |
| 49 | Mut Torsion FoldX | Energetics | 0.64 | 0.63 |
| 50 | Mut Cis bond FoldX | Energetics | 0.63 | 0.59 |
| 51 | WT Cis bond FoldX | Energetics | 0.62 | 0.60 |
| 52 | Mut BackHbond FoldX | Energetics | 0.61 | 0.58 |
| 53 | WT FoldX | Energetics | 0.60 | 0.59 |
| 54 | WT BackHbond FoldX | Energetics | 0.60 | 0.59 |
| 55 | Mut FoldX | Energetics | 0.59 | 0.59 |
| 56 | WT Electro FoldX | Energetics | 0.59 | 0.59 |
| 57 | WT SolvH FoldX | Energetics | 0.58 | 0.57 |
| 58 | WT Entropy SC FoldX | Energetics | 0.58 | 0.59 |
| 59 | WT Ionization | Energetics | 0.58 | 0.58 |
| 60 | Mut SolvH FoldX | Energetics | 0.58 | 0.57 |
| 61 | Mut Entropy SC FoldX | Energetics | 0.57 | 0.60 |
| 62 | Mut Electro FoldX | Energetics | 0.56 | 0.55 |
| 63 | Mut Ionization | Energetics | 0.56 | 0.55 |
| 64 | Mut VdW FoldX | Energetics | 0.55 | 0.57 |
| 65 | WT SideHbond FoldX | Energetics | 0.54 | 0.54 |
| 66 | WT VdW FoldX | Energetics | 0.54 | 0.54 |
| 67 | WT SolvP FoldX | Energetics | 0.54 | 0.54 |
| 68 | Mut SolvP FoldX | Energetics | 0.54 | 0.55 |
| 69 | Mut Backbone_vdwclash FoldX | Energetics | 0.54 | 0.55 |
| 70 | WT VdWclash FoldX | Energetics | 0.53 | 0.53 |
| 71 | WT Backbone_vdwclash FoldX | Energetics | 0.53 | 0.55 |
| 72 | WT Helix Dipole FoldX | Energetics | 0.52 | 0.52 |

| | | | | |
|---|---|---|---|---|
| 73 | Mut SideHbond FoldX | Energetics | 0.52 | 0.53 |
| 74 | Mut VdWclash FoldX | Energetics | 0.50 | 0.50 |
| 75 | Mut Helix Dipole FoldX | Energetics | 0.50 | 0.50 |
| 76 | Protspom Energy | Energetics | 0.33 | 0.32 |
| **Sum** | | | **19.51** | **19.44** |
| 77 | WT Long Range Contact Order (LRCO) | Structural | 2.31 | 2.37 |
| 78 | WT Disulfide Bond | Structural | 1.50 | 1.38 |
| 79 | WT Relative Solvent Accessibility (RSA) | Structural | 1.29 | 1.35 |
| 80 | WT Hydrophobic Buried Core Score (HBCS) | Structural | 0.80 | 0.99 |
| 81 | WT Weighted Contact Number (WCN) | Structural | 0.71 | 0.76 |
| 82 | WT Hydrogen Bond | Structural | 0.63 | 0.61 |
| 83 | Mut HBCS | Structural | 0.57 | 1.06 |
| 84 | Mut Secondary Structure Propensity | Structural | 0.46 | 0.46 |
| 85 | WT Secondary Structure Propensity | Structural | 0.45 | 0.45 |
| 86 | WT weighted hydrophobic contact number (WHbCN) | Structural | 0.45 | 0.46 |
| 87 | WT Hydrophobic Core Score (HCS) | Structural | 0.38 | 0.37 |
| 88 | Mut HCS | Structural | 0.38 | 0.37 |
| 89 | WT Aromatic Cluster Size | Structural | 0.36 | 0.37 |
| 90 | WT Salt bridge | Structural | 0.35 | 0.36 |
| 91 | Mut Hydrogen Bond | Structural | 0.35 | 0.33 |
| 92 | Mut Aromatic Cluster Size | Structural | 0.34 | 0.34 |
| 93 | Mut Saltbridge | Structural | 0.33 | 0.32 |
| **Sum** | | | **11.66** | **12.35** |
| 94 | WT Hydrophobicity Value | Physicochemical | 1.10 | 1.11 |
| 95 | Mut Hydrophobicity Value | Physicochemical | 0.85 | 0.87 |
| 96 | WT Molecular Weight | Physicochemical | 0.65 | 0.65 |
| 97 | Mut Chemical group | Physicochemical | 0.51 | 0.50 |
| 98 | WT Chemical group | Physicochemical | 0.47 | 0.48 |
| 99 | Mut Molecular Weight | Physicochemical | 0.44 | 0.44 |
| **Sum** | | | **4.02** | **4.05** |
| 100 | res_ScanNet Value | Intermolecular Interactions | 0.53 | 0.53 |
| **Sum** | | | **0.53** | **0.53** |

*The structural, intermolecular interactions, evolutionary, energetics, physicochemical, and dynamics features contribute 11.7%, 0.5%, 42.5%, 19.5%, 4% and 21.8% respectively when evaluated on class-stratified 10-fold cross validation. The same features contribute 12.4%, 0.5%, 41.4%, 19.4%, 4% and 22.2% when evaluated on protein-stratified 10-fold validation. The percentage contributions are similar in both cases and do not change the overall picture of which types of features contribute more or less.*

**Table S2.** Performance Evaluation of Rhapsody-2 and its Variants for simple 5-Fold Stratified and Protein-Stratified Cross-Validation on the UnifyPDBFull and UnifyPDBAcceptable Datasets

| # | Method | UnifyPDBFull dataset | | | | UnifyPDBAcceptable dataset | | | |
|---|---|---|---|---|---|---|---|---|---|
| | | Simple 5-Fold CV | | Protein-Stratified 5-Fold CV | | Simple 5-Fold CV | | Protein-Stratified 5-Fold CV | |
| | | **MCC** | **AUROC** | **MCC** | **AUROC** | **MCC** | **AUROC** | **MCC** | **AUROC** |
| 1. | PROVEAN | 0.65 | 0.89 | - | - | 0.63 | 0.89 | - | - |
| 2. | Polyphen-2 | 0.69 | 0.92 | - | - | 0.69 | 0.92 | - | - |
| 3. | PMUT | 0.43 | 0.77 | - | - | 0.47 | 0.80 | - | - |
| 4. | LIST | 0.49 | 0.83 | - | - | 0.51 | 0.84 | - | - |
| 5. | FATHMM | 0.29 | 0.69 | - | - | 0.28 | 0.67 | - | - |
| 6. | EVmutation | 0.69 | 0.78 | - | - | 0.68 | 0.75 | - | - |
| 7. | SPRI | 0.75 | 0.94 | - | - | 0.74 | 0.94 | - | - |
| 8. | *Rhapsody* | 0.73 | 0.92 | - | - | 0.72 | 0.92 | - | - |
| 9. | *Rhapsody-2* | 0.78 | 0.96 | 0.70 | 0.93 | 0.75 | 0.95 | 0.67 | 0.92 |
| 10. | *Rhapsody-2* (red) | 0.74 | 0.94 | 0.69 | 0.92 | 0.72 | 0.93 | 0.66 | 0.91 |
| 11. | *Rhapsody-2* (evo) | 0.71 | 0.92 | 0.65 | 0.91 | 0.69 | 0.92 | 0.64 | 0.91 |
| 12. | *Rhapsody-2* (dyn) | 0.59 | 0.88 | 0.38 | 0.78 | 0.55 | 0.86 | 0.35 | 0.77 |

*#The coverage (the fraction of SAVs out of 9,484 instances in the UnifyPDBAcceptable dataset that could be used for 5-fold cross-validation) of PROVEAN, PolyPhen-2, PMUT, LIST, FATHMM, EVmutation and Rhapsody for UnifyPDBAcceptable were 0.99, 0.99, 0.99, 0.99, 0.99, 0.71, and 0.94 respectively. SPRI and all models of Rhapsody-2 had full coverage. The respective coverages for UnifyPDBFull were 0.99, 0.99, 0.99, 0.99, 0.99, 0.74 and 0.73 respectively, while SPRI and all models of Rhapsody-2 again had full coverage.*

*(-) indicates that there is no data for protein stratified 5-fold cross-validation*

**Table S3.** Distribution of features across neutral and pathogenic SAVs in *Rhapsody-2* SAV-DB (*)

| Feature | Neutral | | | Pathogenic | | | Two-sample t-test p-value |
|---|---|---|---|---|---|---|---|
| | Median | IQR1 | IQR3 | Median | IQR1 | IQR3 | |
| res_MSF (ANM hot) | 0.000 | 0.000 | 0.001 | 0.001 | 0.000 | 0.004 | 0.000e+00 |
| res_MSF_zscore (ANM hot) | 0.002 | 0.000 | 0.022 | 0.036 | 0.009 | 0.113 | 0.000e+00 |
| res_MSF_norm_zscore (ANM hot) | -0.335 | -0.422 | -0.234 | -0.136 | -0.329 | 0.526 | 0.000e+00 |
| res_MSF (GNM all) | 0.470 | 0.148 | 2.068 | 0.136 | 0.075 | 0.412 | 0.000e+00 |
| res_MSF_norm_zscore (GNM hot) | 0.000 | 0.000 | 0.000 | 0.001 | 0.000 | 0.014 | 0.000e+00 |
| res_MSF_norm_zscore (GNM all) | 0.076 | 0.020 | 0.262 | 0.016 | 0.005 | 0.060 | 0.000e+00 |
| res_MSF (GNM soft) | 0.328 | 0.062 | 1.726 | 0.063 | 0.017 | 0.329 | 0.000e+00 |
| res_MSF_norm_zscore (GNM soft) | 0.052 | 0.011 | 0.238 | 0.010 | 0.003 | 0.045 | 0.000e+00 |
| res_MSF_zscore (GNM hot) | -0.198 | -0.212 | -0.185 | -0.185 | -0.207 | -0.100 | 0.000e+00 |
| res_MSF (GNM top5) | 0.203 | 0.040 | 1.176 | 0.045 | 0.011 | 0.246 | 0.000e+00 |
| res_hinge_binary (GNM soft) | 1.000 | 0.000 | 1.000 | 1.000 | 0.000 | 1.000 | 2.291e-33 |
| res_MSF (ANM all) | 142.955 | 12.044 | 1455.995 | 30.360 | 2.251 | 173.736 | 0.000e+00 |
| res_effectiveness (GNM soft) | 0.272 | 0.100 | 0.725 | 0.707 | 0.315 | 1.743 | 0.000e+00 |
| res_zscore_effectiveness (GNM all) | -0.485 | -0.943 | 0.370 | 0.466 | -0.378 | 1.197 | 0.000e+00 |
| res_norm_zscore_effectiveness (GNM all) | 0.234 | 0.081 | 0.479 | 0.467 | 0.253 | 0.712 | 0.000e+00 |
| res_avg_meancorr (GNM soft) | 0.408 | 0.272 | 0.575 | 0.474 | 0.355 | 0.630 | 0.000e+00 |
| res_zscore_sensitivity (ANM top5) | -0.170 | -0.285 | -0.086 | -0.175 | -0.290 | -0.112 | 1.872e-51 |
| res_effectiveness (GNM all) | 0.082 | 0.031 | 0.244 | 0.122 | 0.033 | 0.399 | 1.642e-303 |
| res_norm_zscore_sensitivity (ANM all) | 0.008 | 0.001 | 0.049 | 0.002 | 0.001 | 0.014 | 1.246e-280 |
| res_MSF (ANM soft) | 137.887 | 11.497 | 1374.490 | 30.002 | 2.000 | 171.352 | 0.000e+00 |
| res_effectiveness (ANM all) | 1.007 | 0.103 | 3.952 | 3.269 | 0.711 | 8.519 | 0.000e+00 |
| res_ScanNet | 0.316 | 0.107 | 0.602 | 0.143 | 0.021 | 0.489 | 0.000e+00 |
| WT Mol Wt | 131.000 | 115.000 | 147.000 | 131.000 | 105.000 | 155.000 | 2.633e-04 |
| WT Hydro | -0.800 | -3.500 | 1.800 | -0.700 | -3.500 | 1.900 | 1.150e-05 |
| WT Saltbridge | 0.000 | 0.000 | 0.000 | 0.000 | 0.000 | 0.000 | 6.651e-148 |
| WT Disul | 0.000 | 0.000 | 0.000 | 0.000 | 0.000 | 0.000 | 0.000e+00 |
| WT Hbond | 1.000 | 0.000 | 2.000 | 2.000 | 1.000 | 2.000 | 0.000e+00 |
| WT Aromatic | 0.000 | 0.000 | 0.000 | 0.000 | 0.000 | 0.000 | 9.972e-187 |
| WT HCS | 0.250 | 0.000 | 0.430 | 0.360 | 0.210 | 0.500 | 0.000e+00 |
| WT HBCS | 0.000 | 0.000 | 0.140 | 0.140 | 0.000 | 0.290 | 0.000e+00 |
| WT SS Prop | 0.980 | 0.770 | 1.430 | 1.080 | 0.930 | 1.430 | 8.179e-82 |
| WT FoldX | 953.470 | 490.510 | 2010.500 | 664.880 | 352.280 | 1461.990 | 0.000e+00 |
| Mut Mol Wt | 131.000 | 117.000 | 147.000 | 131.000 | 117.000 | 155.000 | 2.037e-150 |
| Mut Hydro | -0.700 | -3.500 | 2.800 | -0.900 | -3.500 | 2.500 | 6.645e-218 |
| Mut Saltbridge | 0.000 | 0.000 | 0.000 | 0.000 | 0.000 | 0.000 | 3.973e-26 |
| Mut Hbond | 1.000 | 0.000 | 2.000 | 1.000 | 1.000 | 2.000 | 0.000e+00 |
| Mut Aromatic | 0.000 | 0.000 | 0.000 | 0.000 | 0.000 | 0.000 | 4.285e-246 |
| Mut HCS | 0.250 | 0.000 | 0.430 | 0.360 | 0.210 | 0.500 | 0.000e+00 |
| Mut HBCS | 0.000 | 0.000 | 0.140 | 0.140 | 0.000 | 0.290 | 0.000e+00 |
| Mut SS Prop | 0.980 | 0.740 | 1.300 | 0.980 | 0.740 | 1.190 | 4.579e-26 |
| Mut FoldX | 954.720 | 491.455 | 2011.465 | 668.460 | 355.580 | 1467.390 | 0.000e+00 |
| WT RSA | 0.540 | 0.100 | 0.810 | 0.110 | 0.000 | 0.420 | 0.000e+00 |
| WT WCN | 0.670 | 0.380 | 1.040 | 1.080 | 0.690 | 1.410 | 0.000e+00 |
| WT LRCO | 0.180 | 0.000 | 0.570 | 0.400 | 0.250 | 0.500 | 0.000e+00 |
| BLOSSUM (WT, Mut) | 0.000 | -1.000 | 1.000 | -1.000 | -2.000 | 0.000 | 0.000e+00 |
| WT WHbCN | -0.130 | -0.400 | -0.000 | 0.000 | -0.340 | 0.220 | 0.000e+00 |
| WT BackHbond FoldX | -389.100 | -687.720 | -238.895 | -355.700 | -603.040 | -233.200 | 8.909e-37 |
| WT SideHbond FoldX | -84.850 | -135.690 | -49.130 | -69.700 | -127.630 | -46.220 | 2.878e-104 |
| WT VdW FoldX | -685.080 | -1173.930 | -413.980 | -603.160 | -1111.920 | -413.630 | 8.628e-32 |
| WT Electro FoldX | -15.770 | -31.000 | -5.430 | -16.690 | -30.150 | -8.170 | 3.606e-13 |
| WT SolvP FoldX | 989.570 | 609.740 | 1661.750 | 866.700 | 597.770 | 1582.390 | 1.002e-39 |

| | | | | | | | |
|---|---|---|---|---|---|---|---|
| WT SolvH FoldX | -877.980 | -1505.050 | -521.030 | -781.810 | -1374.310 | -519.990 | 1.259e-32 |
| WT VdWclash FoldX | 324.220 | 173.210 | 529.490 | 264.870 | 159.450 | 432.660 | 0.000e+00 |
| WT Torsion FoldX | 82.660 | 38.480 | 236.820 | 58.540 | 27.730 | 123.670 | 0.000e+00 |
| WT Backbone_vdwclash FoldX | 391.240 | 252.300 | 648.790 | 361.560 | 234.510 | 590.140 | 8.973e-24 |
| WT Entropy SC FoldX | 346.390 | 214.870 | 585.370 | 293.560 | 200.080 | 555.800 | 4.205e-71 |
| WT Entropy MC FoldX | 1607.740 | 857.795 | 2418.820 | 1130.370 | 692.490 | 2079.180 | 0.000e+00 |
| WT Helix Dipole FoldX | -5.980 | -12.390 | -2.380 | -6.400 | -12.430 | -2.870 | 3.494e-02 |
| WT Cis bond FoldX | 5.190 | 1.130 | 23.270 | 2.420 | 0.890 | 10.010 | 0.000e+00 |
| WT Disulphide FoldX | 0.000 | -2.200 | 0.000 | 0.000 | -6.640 | 0.000 | 1.359e-147 |
| WT Ionization | 2.110 | 1.170 | 3.320 | 1.710 | 1.020 | 2.800 | 1.177e-273 |
| Mut BackHbond FoldX | -389.120 | -687.765 | -238.855 | -355.660 | -602.990 | -233.140 | 4.733e-37 |
| Mut SideHbond FoldX | -84.820 | -135.690 | -49.130 | -69.700 | -128.130 | -46.120 | 1.174e-106 |
| Mut VdW FoldX | -685.160 | -1173.985 | -413.950 | -603.240 | -1110.600 | -414.060 | 1.025e-31 |
| Mut Electro FoldX | -15.710 | -30.940 | -5.410 | -16.600 | -30.210 | -8.170 | 1.281e-11 |
| Mut SolvP FoldX | 989.750 | 609.550 | 1661.805 | 868.580 | 599.400 | 1583.780 | 1.760e-39 |
| Mut SolvH FoldX | -878.300 | -1504.770 | -521.180 | -781.640 | -1375.460 | -519.290 | 1.455e-32 |
| Mut VdWclash FoldX | 324.540 | 173.850 | 529.870 | 270.580 | 162.470 | 437.350 | 0.000e+00 |
| Mut Torsion FoldX | 82.620 | 38.495 | 237.225 | 58.540 | 27.920 | 123.730 | 0.000e+00 |
| Mut Backbone_vdwclash FoldX | 391.240 | 252.215 | 648.940 | 361.340 | 234.710 | 588.650 | 6.659e-24 |
| Mut Entropy SC FoldX | 346.510 | 214.615 | 585.505 | 293.520 | 200.070 | 557.040 | 3.651e-71 |
| Mut Entropy MC FoldX | 1607.650 | 857.740 | 2418.480 | 1129.970 | 692.560 | 2079.210 | 0.000e+00 |
| Mut Helix Dipole FoldX | -5.980 | -12.380 | -2.380 | -6.380 | -12.490 | -2.870 | 1.869e-02 |
| Mut Cis bond FoldX | 5.190 | 1.130 | 23.275 | 2.420 | 0.850 | 10.010 | 0.000e+00 |
| Mut Disulphide FoldX | 0.000 | -2.200 | 0.000 | 0.000 | -6.610 | 0.000 | 3.583e-143 |
| Mut Ionization | 2.120 | 1.170 | 3.330 | 1.710 | 1.030 | 2.800 | 3.001e-276 |
| WT Chemgroup | 3.000 | 1.000 | 4.000 | 2.000 | 1.000 | 4.000 | 3.094e-04 |
| Mut Chemgroup | 3.000 | 1.000 | 3.000 | 3.000 | 1.000 | 4.000 | 5.386e-109 |
| Protspom Energy | -0.730 | -1.090 | -0.330 | -0.780 | -1.140 | -0.390 | 1.182e-80 |
| WT Seq_PSIC (with gaps) | 1.440 | 0.583 | 2.140 | 2.530 | 1.810 | 3.060 | 0.000e+00 |
| WT PhyChem Group Seq_PSIC (with gaps) | 0.561 | -0.182 | 1.040 | 1.130 | 0.739 | 1.620 | 0.000e+00 |
| WT Size Group Seq_PSIC (with gaps) | 0.488 | -0.257 | 1.010 | 1.280 | 0.853 | 1.640 | 0.000e+00 |
| Mut Seq_PSIC (with gaps) | -0.371 | -1.810 | 0.814 | -2.990 | -4.490 | -1.380 | 0.000e+00 |
| Mut PhyChem Group Seq_PSIC (with gaps) | -0.066 | -1.290 | 0.710 | -1.100 | -3.660 | 0.788 | 0.000e+00 |
| Mut Size Group Seq_PSIC (with gaps) | -0.422 | -1.650 | 0.411 | -1.940 | -4.030 | -0.099 | 0.000e+00 |
| WT-Mut Seq_PSIC (with gaps) | 1.707 | 0.400 | 3.236 | 5.280 | 3.570 | 7.040 | 0.000e+00 |
| WT-Mut PhyChem Group Seq_PSIC (with gaps) | 0.000 | 0.000 | 1.433 | 2.062 | 0.000 | 4.880 | 0.000e+00 |
| WT-Mut Size Group Seq_PSIC (with gaps) | 0.651 | 0.000 | 1.880 | 3.020 | 0.823 | 5.260 | 0.000e+00 |
| WT Seq_Shannon Entropy (with gaps) | 2.120 | 1.270 | 2.830 | 1.280 | 0.685 | 2.040 | 0.000e+00 |
| WT zscore Seq_Shannon Entropy (with gaps) | 0.351 | -0.536 | 1.050 | -0.665 | -1.200 | 0.124 | 0.000e+00 |
| WT PhyChem Group Seq_Shannon Entropy (with gaps) | 1.360 | 0.838 | 1.790 | 0.930 | 0.433 | 1.310 | 0.000e+00 |
| WT PhyChem Group zcore Seq_Shannon Entropy (with gaps) | 0.323 | -0.533 | 1.020 | -0.582 | -1.140 | 0.169 | 0.000e+00 |
| WT Size Group Seq_Shannon Entropy (with gaps) | 1.550 | 1.000 | 1.930 | 1.010 | 0.493 | 1.440 | 0.000e+00 |
| WT Size Group zcore Seq_Shannon Entropy (with gaps) | 0.408 | -0.466 | 1.010 | -0.625 | -1.230 | 0.176 | 0.000e+00 |
| WT Seq_Shannon Entropy (without gaps) | 2.390 | 1.650 | 3.030 | 1.010 | 0.321 | 1.980 | 0.000e+00 |
| WT zscore Seq_Shannon Entropy (without gaps) | 0.577 | -0.133 | 1.080 | -1.010 | -1.540 | -0.098 | 0.000e+00 |
| WT PhyChem Group Seq_Shannon Entropy (without gaps) | 1.230 | 0.742 | 1.600 | 0.455 | 0.098 | 0.998 | 0.000e+00 |
| WT PhyChem Group zcore Seq_Shannon Entropy (without gaps) | 0.526 | -0.284 | 1.070 | -0.939 | -1.370 | -0.007 | 0.000e+00 |

| | | | | | | | |
|---|---|---|---|---|---|---|---|
| WT Size Group Seq_Shannon Entropy (without gaps) | 1.460 | 1.030 | 1.760 | 0.581 | 0.144 | 1.170 | 0.000e+00 |
| WT Size Group zcore Seq_Shannon Entropy (without gaps) | 0.596 | -0.092 | 1.030 | -1.020 | -1.560 | -0.017 | 0.000e+00 |

*(*) "Neutral Median" and "Pathogenic Median" represent the median values of the feature for neutral and pathogenic SAVs, respectively. " IQR1" is the first quartile (25th percentile), while " IQR3" is the third quartile (75th percentile) of the feature, respectively. The "Two-sample t-test p-value" provides the p-value from the two-sample t-test comparing the Neutral and Pathogenic classes. This test assesses whether there is a statistically significant difference between the means of the two classes. It assumes unequal variances (Welch's t-test) and handles missing data with pairwise deletion. A p-value less than 0.05 is typically considered to be statistically significant.*

#### SUPPLEMENTARY FIGURES

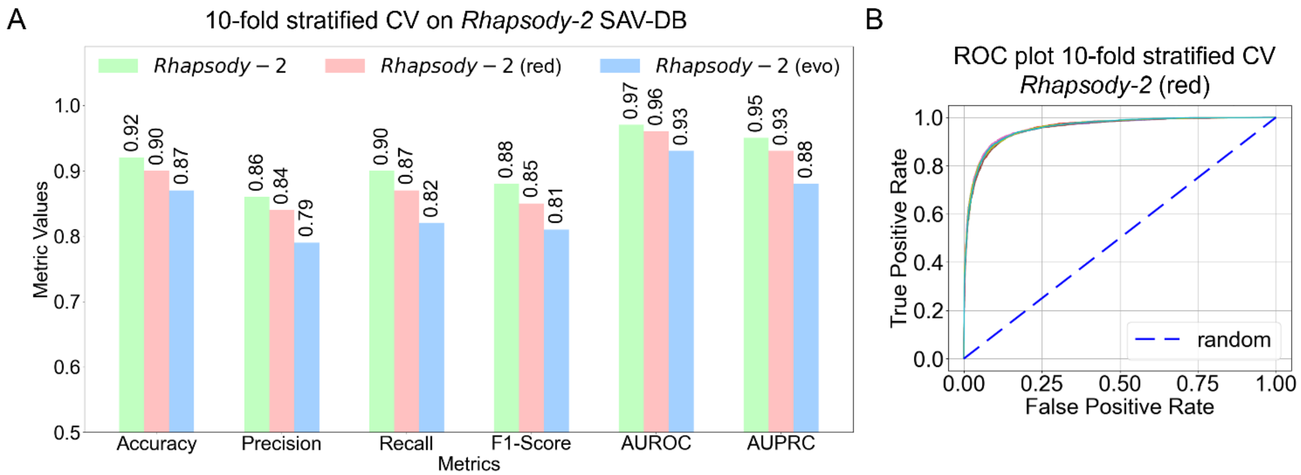

**Figure S1. Performance of *Rhapsody-2* and its reduced model *Rhapsody-2* (red) under 10-fold stratified cross-validation.** (A) Performance metrics of *Rhapsody-2* evaluated using 10-fold stratified cross-validation. (B) ROC curves for the reduced model *Rhapsody-2* (red) under 10-fold stratified cross-validation. The dashed line indicates random behavior.

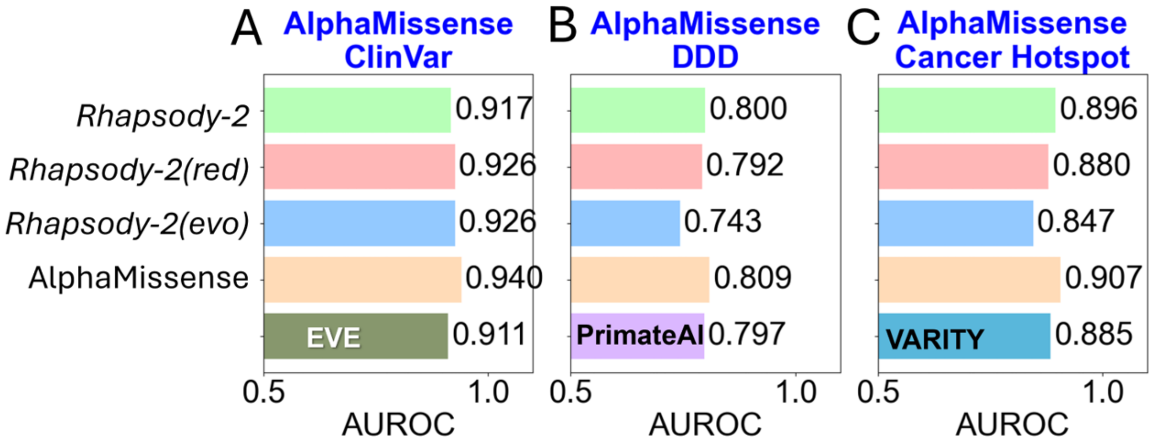

**Figure S2. Rhapsody-2 and its variants exhibit performances comparable to AlphaMissense, EVE, and PrimateAI** when trained on the Rhapsody-2 DB and tested against the AlphaMissense (**A**) ClinVar, (**B**) DDD, and (**C**) Cancer HotSpot datasets. In these tests, while no overlapping SAVs were present between the training and test sets, other variants of the proteins included in the test sets were not removed from the training data.

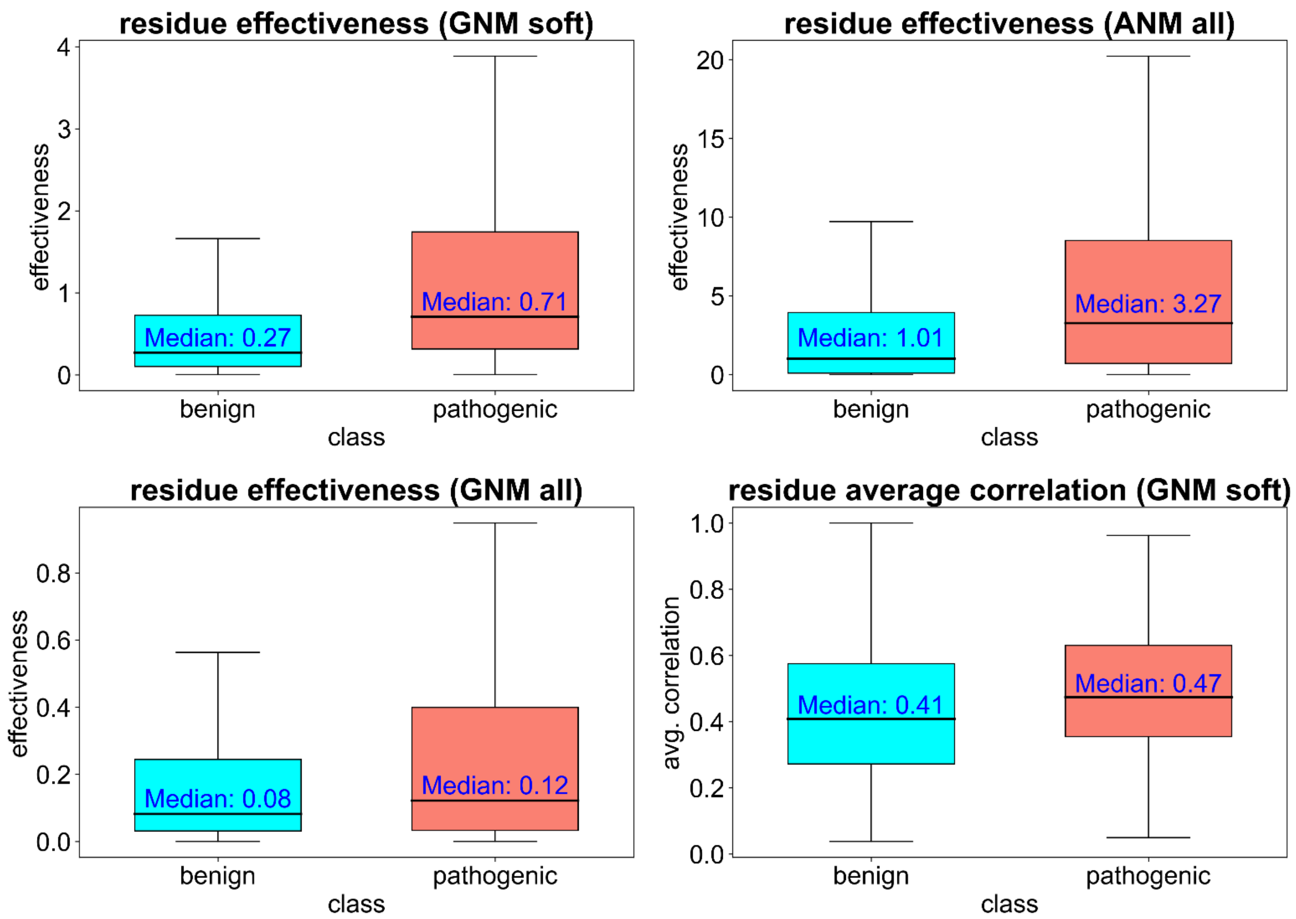

**Figure S3. Distribution of effectiveness and average cross-correlation value for the subsets of pathogenic and neutral/benign mutations.** Allosteric signaling effectiveness computed by the GNM soft modes (*top left*), ANM all modes (*top right*), GNM all modes (*bottom left*), and to a lower extent, the residue correlations based on GNM soft modes (*bottom right*) are examples of dynamic features that help distinguish between neutral and pathogenic SAVs.

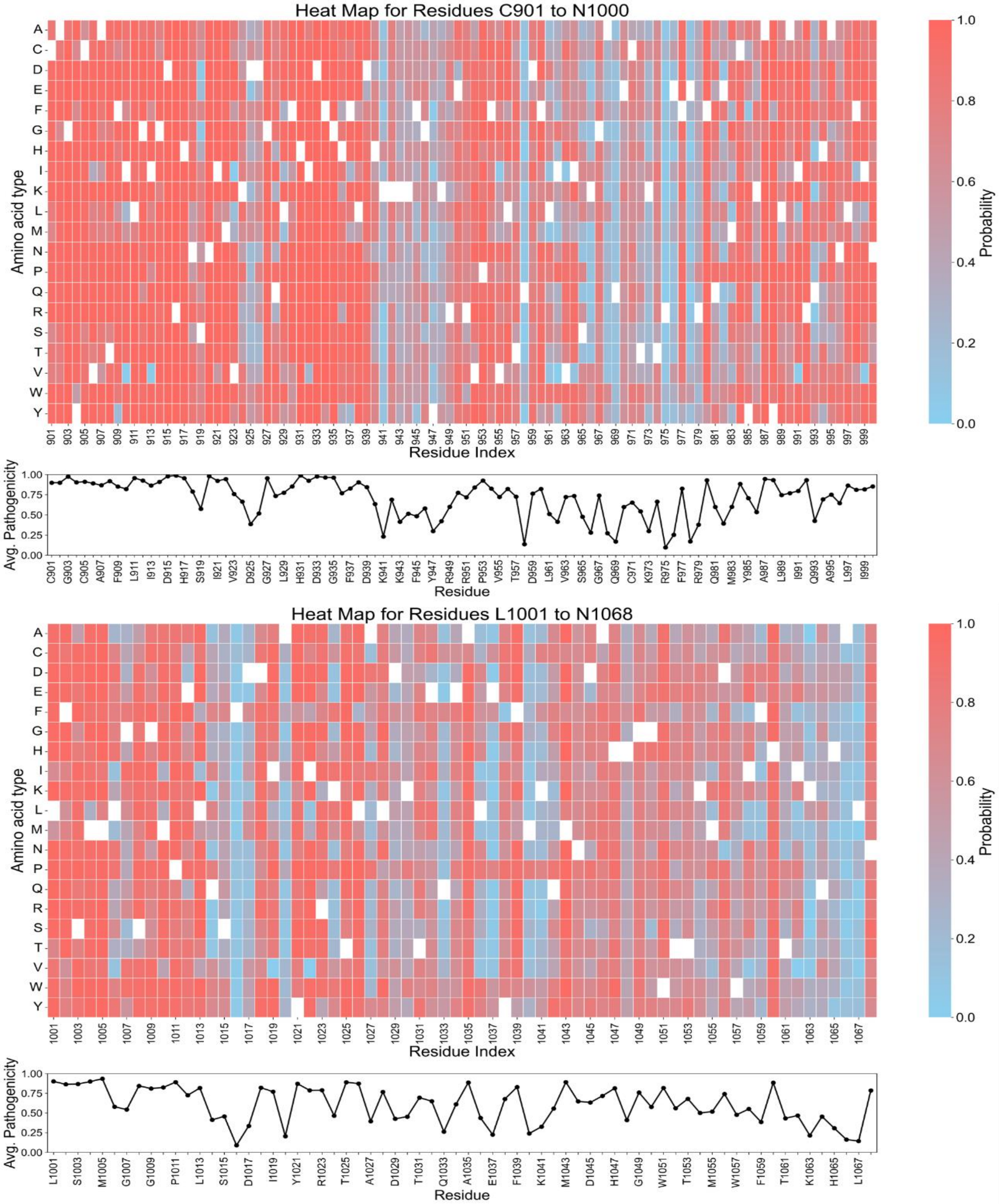

**Figure S4. *In silico* saturation mutagenesis map for PIK3CA generated by *Rhapsody-2* (*red*).** The panels show the heatmaps for all possible mutations (*ordinate*) at positions C901-N1000 (*top*) and L1001-A1068 (*bottom*), as predicted by *Rhapsody-2*(*red*). Computations were performed for the complete structure (of 1068 amino acids) using the AlphaFold2 predicted structural model for PIK3CA (AF-P42336-F1-model_v4.pdb). Results are displayed for the N-terminal portion (between C901 and A1068) to focus on four SAVS, T1025A, Y1021H, Y1021C, and H1047R, each reported (in ClinVar) to be pathogenic. The map entries are color-coded based on predicted probability of pathogenicity, from 0 (*blue*, neutral) to 1 (*red*, highly pathogenic). The entries corresponding to WT amino acids are in *white*. The curve under the heatmaps refers to the average pathogenicity score evaluated for all possible mutations of that particular residue (averaged over all entries in the corresponding column of the map). The ribbon diagram in **Figure 6A** is color-coded by these average pathogenicity values for each residue (M1-N1068).

#### SUPPLEMENTARY REFERENCES

1. M. J. Landrum *et al.*, ClinVar: improving access to variant interpretations and supporting evidence. *Nucleic Acids Res* **46**, D1062-d1067 (2018).
2. M. Varadi *et al.*, AlphaFold Protein Structure Database: massively expanding the structural coverage of protein-sequence space with high-accuracy models. *Nucleic Acids Res* **50**, D439-d444 (2022).
3. B. Wang *et al.*, Structure-based pathogenicity relationship identifier for predicting effects of single missense variants and discovery of higher-order cancer susceptibility clusters of mutations. *Brief Bioinform* **24** (2023).
4. J. Cheng *et al.*, Accurate proteome-wide missense variant effect prediction with AlphaMissense. *Science* **381**, eadg7492 (2023).
5. L. Sundaram *et al.*, Predicting the clinical impact of human mutation with deep neural networks. *Nat Genet* **50**, 1161-1170 (2018).
6. M. T. Chang *et al.*, Accelerating Discovery of Functional Mutant Alleles in Cancer. *Cancer Discov* **8**, 174-183 (2018).
7. F. E. Dewey *et al.*, Distribution and clinical impact of functional variants in 50,726 whole-exome sequences from the DiscovEHR study. *Science* **354** (2016).
8. J. Kyte, R. F. Doolittle, A simple method for displaying the hydropathic character of a protein. *J Mol Biol* **157**, 105-132 (1982).
9. P. Y. Chou, G. D. Fasman, Conformational parameters for amino acids in helical, beta-sheet, and random coil regions calculated from proteins. *Biochemistry* **13**, 211-222 (1974).
10. Q. Wang, A. A. Canutescu, R. L. Dunbrack, Jr., SCWRL and MolIDE: computer programs for side-chain conformation prediction and homology modeling. *Nat Protoc* **3**, 1832-1847 (2008).
11. A. Banerjee, P. Mitra, Estimating the Effect of Single-Point Mutations on Protein Thermodynamic Stability and Analyzing the Mutation Landscape of the p53 Protein. *J Chem Inf Model* **60**, 3315-3323 (2020).
12. M. Heinig, D. Frishman, STRIDE: a web server for secondary structure assignment from known atomic coordinates of proteins. *Nucleic Acids Res* **32**, W500-502 (2004).
13. I. K. McDonald, J. M. Thornton, Satisfying hydrogen bonding potential in proteins. *J Mol Biol* **238**, 777-793 (1994).
14. J. Tubiana, D. Schneidman-Duhovny, H. J. Wolfson, ScanNet: an interpretable geometric deep learning model for structure-based protein binding site prediction. *Nat Methods* **19**, 730-739 (2022).
15. S. Zhang *et al.*, *ProDy 2.0*: increased scale and scope after 10 years of protein dynamics modelling with Python. *Bioinformatics* **37**, 3657-3659 (2021).
16. I. Bahar, A. R. Atilgan, B. Erman, Direct evaluation of thermal fluctuations in proteins using a single-parameter harmonic potential. *Fold Des* **2**, 173-181 (1997).
17. A. R. Atilgan *et al.*, Anisotropy of fluctuation dynamics of proteins with an elastic network model. *Biophys J* **80**, 505-515 (2001).
18. I. J. General *et al.*, ATPase subdomain IA is a mediator of interdomain allostery in Hsp70 molecular chaperones. *PLoS Comput Biol* **10**, e1003624 (2014).
19. C. Atilgan, A. R. Atilgan, Perturbation-response scanning reveals ligand entry-exit mechanisms of ferric binding protein. *PLoS Comput Biol* **5**, e1000544 (2009).
20. J. Schymkowitz *et al.*, The FoldX web server: an online force field. *Nucleic Acids Res* **33**, W382-388 (2005).
21. B. Buchfink, K. Reuter, H. G. Drost, Sensitive protein alignments at tree-of-life scale using DIAMOND. *Nat Methods* **18**, 366-368 (2021).

22. S. R. Sunyaev *et al.*, PSIC: profile extraction from sequence alignments with position-specific counts of independent observations. *Protein engineering* **12**, 387-394 (1999).
23. S. Henikoff, J. G. Henikoff, Amino acid substitution matrices from protein blocks. *Proc Natl Acad Sci U S A* **89**, 10915-10919 (1992).
24. T. Chen, C. Guestrin (2016) Xgboost: A scalable tree boosting system. in *Proceedings Of The 22nd Acm Sigkdd International Conference on Knowledge Discovery and Data Mining,* pp 785-794.
